# The brain cortical similarity network: Development and sensitivity to early life stress in a rat model

**DOI:** 10.1101/2024.12.20.629759

**Authors:** Rachel L. Smith, Stephen J. Sawiak, Lena Dorfschmidt, Ethan G. Dutcher, Jolyon A. Jones, Joel D. Hahn, Olaf Sporns, Larry W. Swanson, Paul A. Taylor, Daniel R. Glen, Jeffrey W. Dalley, Francis J. McMahon, Armin Raznahan, Petra E. Vértes, Edward T. Bullmore

**Affiliations:** Department of Psychiatry, University of Cambridge, Cambridge, CB2 0SZ, UK; Human Genetics Branch, National Institute of Mental Health, Bethesda, MD, USA 20892; Behavioural and Clinical Neuroscience Institute, University of Cambridge, Downing Site, Cambridge, UK; Department of Physiology, Development and Neuroscience, University of Cambridge, Cambridge, CB2 3EL, UK; Department of Psychology, University of Cambridge, Cambridge, CB2 3EB, UK; Department of Biological Sciences, University of Southern California, Los Angeles, CA, USA 90089; Indiana University Network Science Institute, Indiana University, Bloomington, IN, USA 47405; Department of Psychological and Brain Sciences, Indiana University, Bloomington, IN, USA 47405; Scientific and Statistical Computing Core, National Institute of Mental Health, NIH, Bethesda, MD, USA 20892

**Keywords:** connectome, architectome, micro-structural MRI, translational neuroscience, structural similarity

## Abstract

Understanding how early life experiences shape brain network development is a key challenge in the neuroscience of mental health disorders. To address this, we used magnetic resonance imaging (MRI) similarity network analysis to study the effects of stress in the rat, an important animal model in neuropsychiatry. We measured magnetization transfer ratio (MTR) at each of 53 distinct cortical areas and estimated a cortical similarity network for each individual scan, in two independent experimental datasets. We first characterized normative network development in rats scanned repeatedly between postnatal days 20 (weanling) and 290 (mid-adulthood) (N=47), and then contrasted these findings with a cohort exposed to early life stress in the form of repeated maternal separation (RMS, N=40). The normative rat cortical similarity network exhibited biologically meaningful organization, aligning with prior cytoarchitectonic and tract-tracing data, and displayed complex topological features, including rich club organization. During postnatal and adolescent development, brain regions became more similar, including an early phase of fronto-hippocampal convergence. Early increases in inter-areal similarity were reversed in a later phase of fronto-hippocampal divergence in mid-adulthood. RMS exposure altered inter-areal similarity, especially between frontal and parahippocampal regions, that were also most active developmentally and in aging. Our results reveal how normative cortical network changes in the developing brain are influenced by early life stress. These findings suggest a new translational framework for elucidating how environmental risk factors lead to atypical development of cortical networks.

## Introduction

Altered trajectories of brain development are a major risk factor for psychiatric and neurodevelopmental disorders (Sánchez et al., 2001; Paus et al., 2008; Lupien et al., 2009). However, the biological mechanisms that govern brain development—and how they are shaped by life experiences—remain incompletely understood. Tools that capture dynamic, systems-level changes in neuroarchitecture can advance our insight into these processes. Network models of macroscale brain organization, particularly those constructed using noninvasive neuroimaging, provide a key framework for characterizing distributed brain structure and function (Bullmore and Sporns, 2009, 2012; van den Heuvel and Sporns, 2011, 2013; Fornito et al., 2015). By quantifying inter-regional relationships, these models reveal the organization and maturation of brain systems across development and in response to experience.

Structural similarity analysis is an emerging approach for constructing individualized brain networks using magnetic resonance imaging (MRI) (Li et al., 2017; Seidlitz et al., 2018; Sebenius et al., 2023). These methods estimate inter-regional similarity using MRI-derived morphometric features (Sebenius et al., 2024), resulting in networks in which nodes represent cortical areas and edges reflect the strength of structural similarity between them. A recent implementation, morphometric inverse divergence (MIND), quantifies similarity between voxel- or vertex-level distributions of MRI features (Sebenius et al., 2023). MIND and related networks are reproducible (Seidlitz et al., 2018; Sebenius et al., 2023), developmentally sensitive (Fenchel et al., 2020; Ruan et al., 2023; Sebenius et al., 2023; Wu et al., 2023; Dorfschmidt et al., 2024), and responsive to environmental exposures (Tian et al., 2021; González-García et al., 2023; Xiao et al., 2023; Hettwer et al., 2024). MIND networks also show clinical relevance (Mahjoub et al., 2018; Homan et al., 2019; Li et al., 2019; Lisowska and Rekik, 2019; Zhang et al., 2020; Cao et al., 2023), significant heritability (Sebenius et al., 2023), and associations with gene expression (Morgan et al., 2019; Seidlitz et al., 2020).

Despite these advantages, structural similarity networks have been almost exclusively studied in humans, limiting their utility for testing causal mechanisms. Animal models are essential for addressing this gap, enabling experimental manipulation and brain tissue access. Rats in particular offer distinct advantages, including suitability for high-throughput experiments and ethologically rich, human-relevant behaviors (Bryda, 2013; Ellenbroek and Youn, 2016). Despite this, network representations of the rat brain remain limited. Existing data, primarily derived from expertly curated tract-tracing studies (Swanson et al., 2017, 2019, 2020, 2022, 2024a, 2024b), suggest that the rat connectome exhibits complex topology (Swanson et al., 2024b) akin to that in humans (Bullmore and Sporns, 2009) and mice (Rubinov et al., 2015). However, these data represent the composite rat brain and thus do not support individual-level analyses.

Here, we extend the MIND framework to rats, enabling construction of individual-level cortical similarity networks from in vivo MRI and providing a tool to examine how early life environments shape brain organization. We focus on magnetization transfer ratio (MTR), a proxy for cortical myelination (Mancini et al., 2020)—a feature that is both developmentally dynamic (Hamano et al., 1998; Downes and Mullins, 2014; Mengler et al., 2014) and environmentally sensitive in rats (Bass et al., 1970; Krigman and Hogan, 1976; Breton et al., 2021; Long et al., 2021; Han et al., 2022; Abraham et al., 2023), but underexplored in rat MRI studies (Bai et al., 1996).

We applied this method to two rat neuroimaging datasets: a longitudinal normative developmental cohort of males scanned across postnatal development (**Fig. 1A**), and an experimental stress cohort of males and females exposed to repeated maternal separation (RMS) as a model of early life stress (**Fig. 1B**). We demonstrate that MIND networks reflect biologically meaningful features of cortical organization that track age- and experience-related changes in network architecture. These findings offer a novel approach for linking early life experiences to brain network development.

**Figure 1.**
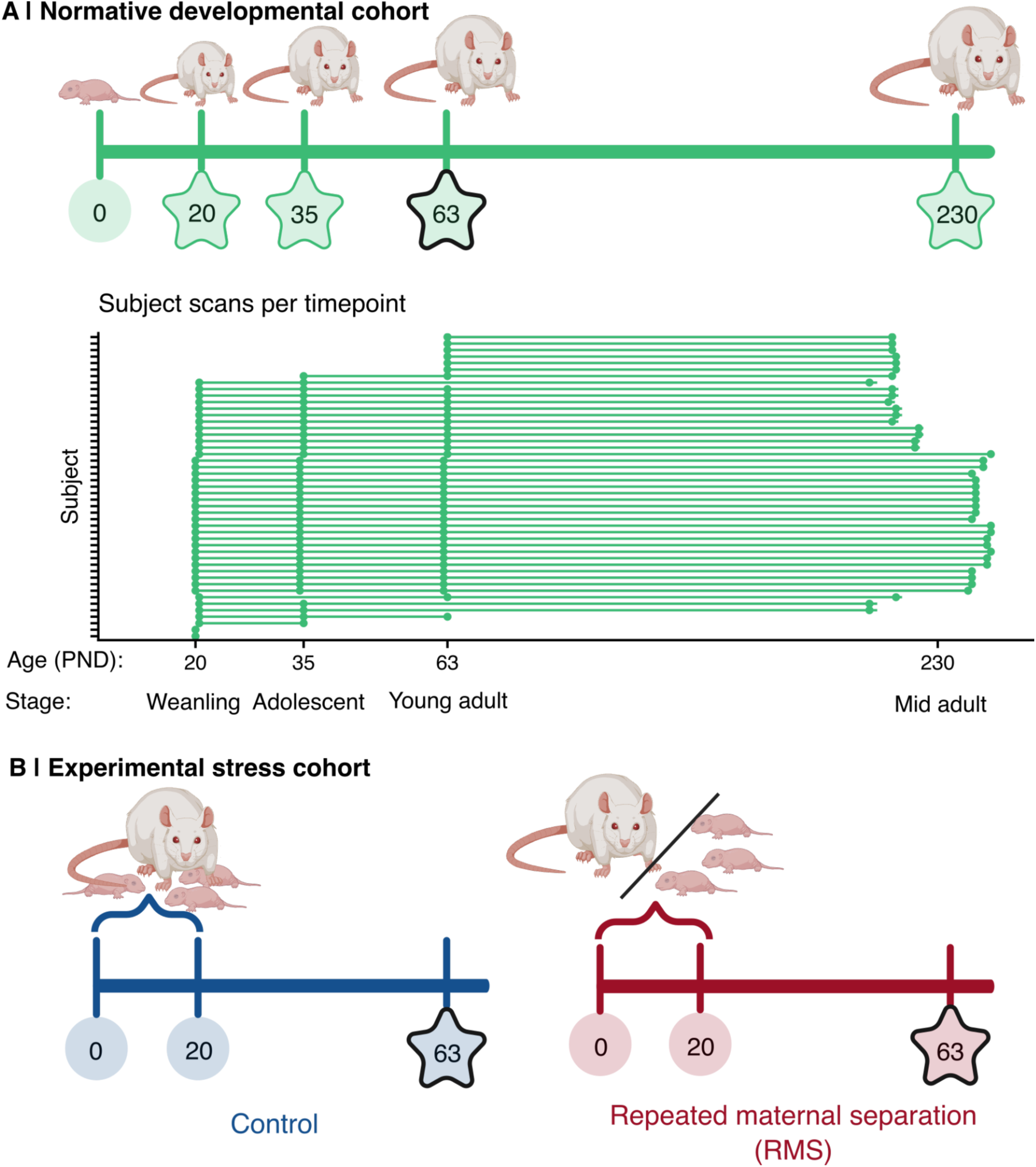
Two independent experimental cohorts to assess reliability and validity of rat MRI similarity networks as measures of developmental and stress-related changes in cortical microstructural networks. **A)** Top: Study design for the normative developmental cohort: N=47 male Lister Hooded rats were reared normally and had brain MRI scanning at PND 20 (N=40), PND 35 (N=38), PND 63 (N=42), and within a few days of PND 230 (N=43). All MRI timepoints are indicated by a star; PND 63 is outlined in black, as it marks the timepoint used to construct the normative network. Bottom: Scans per subject. Each tick on the y-axis represents a subject, while the x-axis shows age. Each point represents a scan; repeated scans of the same rat are connected by a line. **B)** Study design for the experimental stress cohorts. N=21 Lister Hooded pups were stressed by repeated maternal separation (RMS), i.e., for 1 hour/day every day from PND 0 to PND 20, pre-weaning pups were separated from their dam. A control group of N=19 pups was reared normally. All animals completed MRI scanning at PND 63 as young adults. As with panel A, MRI timepoints are indicated by stars, and the primary comparison timepoint (PND 63) is outlined in black. Figure created in https://BioRender.com.

## Materials and Methods

### Experimental design

Pre-existing structural MRI data from two independent cohorts, published in (Jones et al., 2024) and (Dutcher et al., 2023), were used to assess network-level changes that occur during development and in response to stress:

#### 1. Male normative developmental cohort (Jones et al., 2024) (Fig. 1A)

Male Lister Hooded rats (N=47) were kept on a reverse light/dark cycle with red light on from 7:30am - 7:30pm and white light for the other half of the daily cycle. Rats underwent MRI scanning and weaning on postnatal day (PND) 20 or 21 (N=40; here referred to as PND 20). Rats were scanned again at PND 35 (N=38), PND 63 (N=42), and once between PND 212-244 (N=43; here referred to as PND 230). **Figure 1** shows the number of observations per animal in the male normative developmental cohort (hence referred to as the normative developmental cohort), with 32 animals (68% of total) having scans at each of the four timepoints from PND 20 (post-weaning) to PND 230 (aging adult). Experiments were carried out in accordance with the (U.K Animals) Scientific Procedures Act (1986) under UK Home Office project licenses (PPL 70/7587 & PPL 70/8072) and were approved by the University of Cambridge Ethics Committee.

#### 2. Experimental stress cohort (Dutcher et al., 2023) (Fig. 1B)

Pregnant Lister Hooded rats (N=14) were purchased from Envigo (Blackthorn, UK). Litters were delivered by spontaneous partum on gestational days 22-24. Within three days of birth, litter size was adjusted to 4-6 pups, with each litter consisting of two female and two male pups (except for one litter with four males and two females). If two litters were born within 24 hours of one another (the case for 10 litters in total), pups were mixed between the litters. After litter size adjustment, litters were allocated alternately by birth time to either the repeated maternal separation condition (RMS; N=30 pups: 14 female, 16 male) or the control condition (N=28 pups: 14 female, 14 male). PND 0 was defined as the day of delivery. Body weight was measured weekly starting at PND 20. Lights were on from 21:00 to 09:00. Experiments were conducted on Project License PA9FBFA9F, in accordance with the UK Animals (Scientific Procedures) Act 1986 Amendment Regulations 2012, the EU legislation on the protection of animals used for scientific purposes (Directive 2010/63/EU), and the GSK Policy on the Care, Welfare and Treatment of Animal, following ethical review by the University of Cambridge Animal Welfare and Ethical Review Body (AWERB).

From PND 5-19 (inclusive), pups from RMS litters were separated from their dam for 6 hours a day, starting between 11:00am-12:30pm. During separation, dams remained in their home cages while pups were taken to a different room and placed together inside a ventilated cabinet. One centimeter of bedding was provided and the temperature at the surface of the bedding was kept between 30 °C and 35 °C through warming of the air and use of an electric heat pad. Control pups were subject only to normal animal facility rearing. Following PND 20, pups from both groups were weaned and housed in same-sex pairs at PND 20 and left undisturbed until early adulthood except for weighing and once-weekly cage changes. All animals then underwent MRI scanning at PND 63.

### MRI acquisition

For both cohorts, high-resolution MRI was performed on a 9.4T horizontal bore MRI system (Bruker BioSpec 94/20; Bruker Ltd.). Images were acquired under isoflurane anesthesia using the manufacturer-supplied rat brain array coil with the rat in a prone position. Structural images were obtained based on a 3D multi-gradient echo sequence (TR/TE 25/2.4 ms with 6 echo images spaced by 2.1 ms, flip angle 6° with RF spoiling of 117°). The field of view was 30.72 × 25.6 × 20.48 mm3 with a matrix of 192 × 160 × 160 yielding isotropic resolution of 160 μm with a total scan time of 6 min 36 sec with zero-filling acceleration (25% in the readout direction; 20% in each phase encoding direction). Magnetization transfer pulses (10 μT, 2 kHz off-resonance) were applied within each repetition to enhance gray-white matter contrast. Post-reconstruction, images from each echo were averaged after weighting each by its mean signal.

Throughout all scanning procedures, rats were anesthetized with isoflurane (1-2% in 1 L/min O2: air 1:4). Respiratory rate, oxygen saturation and pulse rate (SA Instruments; Stony Brook, NY) were measured with anesthetic dose rates adjusted to ensure readings remained within a physiological range. Body temperature was measured and regulated with a rectal probe and heated water system to 36-37°C.

### Image registration

Structural MRI image preprocessing was performed using the AFNI software package version AFNI_24.2.03 (Cox, 1996). Briefly, magnetization transfer (MT) images for each rat were first deobliqued, spatially oriented, and translated to have spatial overlap with the Waxholm Space reference template (WHS) (Kleven et al., 2023). This space was selected due to its alignment with gold-standard histological rat brain atlases (Paxinos and Watson, 2006; Swanson, 2018) and the atlas’s parcellation granularity (N=222 regions). The @animal_warper command (Jung et al., 2021) was then used to nonlinearly align all MT images to the WHS template. Because some input datasets were notably smaller than the WHS standard template, the -init_scale option was added and used to increase the search space of the registration algorithm, scaled to the relative size ratio of the input scan to the template. Quality control image outputs for each scan (produced by @animal_warper) were then manually reviewed; for any images that were not successfully registered, @animal_warper was run again with the -init_scale parameter altered to better approximate the size ratio. All image registration scripts are available with this publication.

### Magnetization transfer ratio calculation

Magnetization transfer ratio (MTR) was calculated on a per-voxel basis for each scan in native space. First, native scans were scaled according to the receiver gain (RG) parameter used in the scan acquisition protocols. The scaled MT and proton density (PD)-weighted scans in native space were then used to calculate MTR according to the equation (PD - MT) / PD. Both steps were executed using 3dcalc in AFNI (Cox, 1996).

Manual evaluation of MTR data quality resulted in the exclusion of 7 scans, typically because of motion artifacts and particularly at the earlier timepoints. Example MT images and their quality control images are shown in **Figure S1**. The analyzable MRI datasets available following preprocessing and quality control of the two cohorts are summarized in **Table 1**.

**Table 1:** Number of MRI scans included in the study analyses following quality control. *N* represents the number of unique individuals scanned at each timepoint within each cohort.

|  | postnatal days (PND) | *N* |
| --- | --- | --- |
| **Normative developmental cohort** |  |  |
|  | 20 | 40 |
|  | 35 | 38 |
|  | 63 | 41 |
|  | 230 | 43 |
| **Experimental stress cohort** |  |  |
| Repeated maternal separation (RMS) | 63 | 21 |
| Normal maternal rearing (control) | 63 | 19 |

### Morphometric INverse Divergence (MIND) network calculation

MIND networks estimate structural similarity from MRI data (Sebenius et al., 2023). Briefly, cortical regions are represented by a distribution of structural MRI features sampled at many points within the region, in this case, at each voxel. The MIND similarity between each pair of regions is then calculated using the Kullback-Leibler (KL) divergence between their feature distributions.

#### Input data generation

Each MTR image was aligned with the WHS atlas in the native space of the respective MT scan (the scan with the highest contrast), so that each voxel in the MTR scan was labeled with a region of interest based on voxel assignment output from the registration pipeline. For each scan, a two-column CSV was generated, in which the first column “Label” was the region of interest, and the second column (“MTR”) was the value for the corresponding voxel in the MTR scan. For cortical MIND calculation, the input CSV was filtered to contain only voxels belonging to regions under the “Cerebral cortex” hierarchical level of the WHS atlas (which excludes olfactory bulb regions).

#### MIND network construction

MIND networks were constructed for each scan by calculating the KL divergence between pairwise combinations of regional MTR profiles using the MIND toolkit (https://github.com/isebenius/MIND). The MIND algorithm used can be sensitive to differences in number of datapoints. To balance this, we estimated KL divergence for each pair of regions by estimating the same number of voxels (5000) from each region, regardless of its size.

#### MIND network phenotypes

In network neuroscience, topography refers to the spatial layout of brain regions, while topology refers to how those regions are interconnected. Here, we focus on the topological structure of the normative cortical MIND network.

Edge weight and nodal strength (or weighted degree) were considered features of interest for downstream analyses. Edge weights were calculated as w = 1 / (1 + KL), per the MIND toolkit (Sebenius et al., 2023). Nodal strength was calculated as the sum of all edge weights for a given region; hubs were defined as the 10 regions with the highest nodal strength. The rich club coefficient (Φ), a network measure that quantifies the tendency of hubs to be more densely interconnected compared to lower-strength nodes (van den Heuvel and Sporns, 2011; Fornito et al., 2016a), was calculated for the normative network as follows:

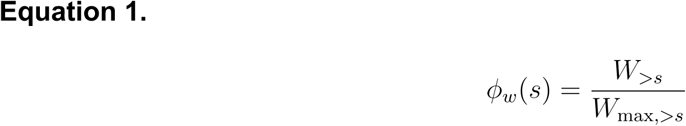

where

- *W*_>*s*_ = sum of edge weights among nodes with strength greater than *s* (i.e., hubs)
- *W*_max, >*s*_ = sum of the strongest edges in the network

### Edge distance calculation

The midline of the WHS atlas was determined using the AFNI function 3dcalc to separate the left and right hemispheres. To approximate the center of each WHS region of interest, the AFNI function 3dCM -Icent (Cox, 1996) was then used. The Euclidean distance between pairwise regional centers was then calculated as their edge length.

### Rat atlas mapping

We mapped the WHS atlas into Brain Maps 4.0 (BM4) atlas space (Swanson, 2018), Zilles atlas space (Zilles, 2012), and Allen Mouse Brain Atlas space (AMBA) (Lein et al., 2007) to compare the MIND similarity network to cortical tract-tracing data (Swanson et al., 2017, 2024b), cortical type (García-Cabezas et al., 2023), and mouse spatial transcriptomic expression (Yao et al., 2023), respectively. We did so first by aligning cortical regions based on nomenclature. However, not all regions maintained consistent nomenclature across atlases, so we also visually inspected anatomical alignment of regions using the WHS EBRAINS online resource (https://www.ebrains.eu/tools/rat-brain), BM4 atlas maps (https://sites.google.com/view/the-neurome-project/brain-maps), Zilles atlas cortical maps in stereotaxic coordinates (available for download at https://link.springer.com/book/10.1007/978-3-642-70573-1), and AMBA online interactive atlas viewer (http://atlas.brain-map.org/atlas?atlas=1). Briefly, we identified the position of each region in the reference (BM4, Zilles, or AMBA) atlas maps, then panned through the WHS atlas using the EBRAINS tool to approximate the same coronal slice and identify what region most closely aligned with the reference anatomical position. We also considered relative positioning of surrounding regions to determine this alignment. All atlas mappings are provided as resources in **Tables S2A-C**.

In this work, the BM4 atlas was used as the reference space for the tract-tracing comparison, the Zilles atlas was used as the reference space for the cortical type comparison, and the AMBA atlas provided reference space for the mouse transcriptomics comparison. If multiple WHS atlas subdivisions comprised a single reference atlas region, the median across these subdivisions was taken as the MIND edge weight for that region.

### Tract-tracing Jaccard index calculation

To convert the cortical tract-tracing matrix (Swanson et al., 2017, 2024b) into a similarity network, the Jaccard index between each pairwise combination of regions was calculated. The set for a given region *a* was defined as the ordinal tract-tracing weight with each other region. The Jaccard index *J* between two regions *a, b* was then calculated as the intersection of their sets divided by the union as follows:

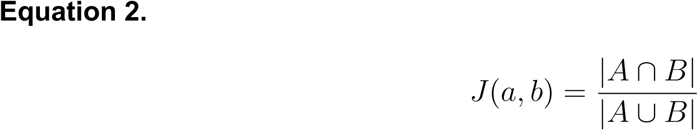

### Cortex type network thresholding

WHS cortical regions were categorized by their cortical type using the data presented in (García-Cabezas et al., 2023) and grouped according to whether they are part of the archicortical allocortex or mesocortex (agranular or dysgranular subdivisions). Paleocortical and eulaminate regions were excluded from this analysis due to the very small number of constituent regions defined by the atlas. Each MIND edge was then defined as ‘intra-class’ or ‘inter-class’ based on whether both regions were part of the same cortex type or not, respectively. To assess the extent to which top-weighted MIND edges consisted of two regions within the same cortex type, the normative MIND network was thresholded across densities (comprising 0-10% of top-weighted edges), and the percentage of intra-class edges was calculated.

### Statistical analysis

#### Null network generation

Ten thousand null networks were generated to assess whether normative MIND network alignment with tract-tracing similarity and cortical type were greater than would be expected by chance. To do so, all MIND network edges were classified into three evenly sized bins based on distance: proximal, intermediate, and distal. Edge weights within each bin were then reshuffled to generate a null network that preserved distance structure. The tract-tracing and cortical type analyses were repeated for each null network to generate a null distribution for comparison of each analysis.

#### Developmental slope

Changes that occurred in edge weight (*w*) and nodal strength (*s*) during normative development (Δw_dev_; Δs_dev_) and aging (Δw_age_; Δs_age_) were quantified by calculating the linear gradient or slope of age-related change in each period. Early development was defined as PND 20 to PND 35, as the highest proportion of change occurred here, and aging was defined as PND 63 to PND 230. For each region, a linear mixed effects model was fit with the normalized strength as the dependent variable, continuous age and total brain volume as fixed effects, and each individual rat as a random effect:

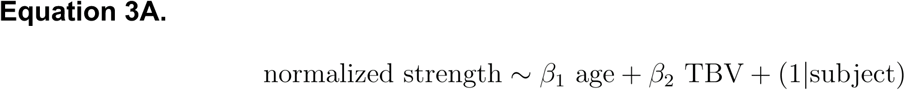

Edge-level analyses were run using coarse-grained systems labels for interpretability. In this case, ROI-level edges were also included as a random effect:

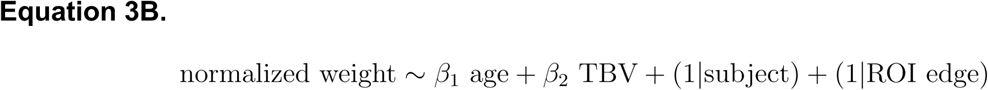

The coefficient for age, β_1_, was estimated as the slope for that region within the given period. The effect size (Δw; Δs) was used to plot and compare between early development and aging epochs.

#### RMS-control effect size

Case-control analysis at PND 63 was used to identify changes that occurred in response to RMS and were measurable in young adulthood. For each region, the following model was used:

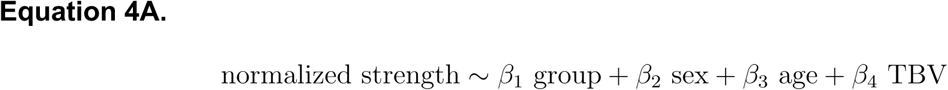

As with development, case-control edge effects were determined at the systems-level, with ROI-level edges included as a random effect:

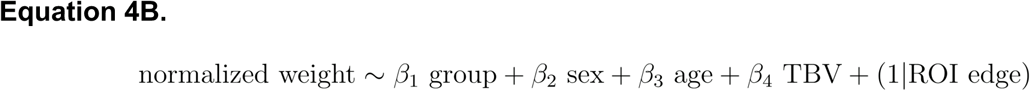

The effect size associated with the group term was estimated as the actual case-control effect size for a given region or edge. Then, for each region and edge, 1000 permutations were run in which the group label was randomly sampled, and the case-control effect size was estimated under the null hypothesis. The *Z*-score for each actual effect size in this permutation distribution was calculated; any region with absolute value *Z*-score > 1.96 (*P*=0.05) was considered significant.

#### Relating developmental changes and case-control stress effects

To characterize the relationship between RMS case-control effects and normative developmental changes, Pearson’s correlation was run on Δw_dev_ and Δw_age_ vs edge-level PND 63 case-control effect size. To assess the extent to which the strength of this relationship was greater than expected under the null hypothesis, 10000 permutations were run, in which the edge assignment of RMS effect size was randomly resampled and again correlated with Δw_dev_ and Δw_age_. The *Z*-score of the actual Pearson’s *r* was determined, and the *P*-value was calculated as 1 - (proportion of permuted correlations that were smaller than the observed correlation).

Across all statistical analyses, *P* values were corrected for multiple comparisons using the Benjamini-Hochberg (BH) procedure (Benjamini and Hochberg, 1995).

### Rat brain plot construction

#### Flatmap rendering

A flatmap is a two-dimensional representation of the brain that facilitates visualization of spatial relationships between regions (Herrick, 1948; Swanson, 2000). Flatmaps are particularly useful for displaying cortical organization and topographic continuity, avoiding distortions introduced by sulcal and gyral folding in three-dimensional renderings (Swanson, 2000). While flatmapping techniques have been used elsewhere in comparative neuroanatomy and cortical cartography (Herrick, 1948; Nauta and Karten, 1970), Swanson and colleagues were the first to systematically adapt this approach to the rat brain (Swanson, 1992; Hahn et al., 2021; Hahn and Duckworth, 2023). In this study, rat brain flatmap plots were derived from the work of (Hahn et al., 2021; Hahn and Duckworth, 2023).

The SVG file provided by (Hahn and Duckworth, 2023) was first converted to well-known text (WKT) using code from https://github.com/davidmcclure/svg-to-wkt. The WKT was then parsed into geometry objects in R using the package sf version 1.0.17 (Pebesma, 2018). Flatmaps represent a practical and underutilized tool for visualizing architectonic variation and similarity in the rat cortex, particularly in connectomic studies. To facilitate broader use, we provide all flatmap construction resources.

#### Anatomical rendering

Rat brain maps were also visualized using an anatomical rendering that preserved 3-dimensional brain structure. Nifti files representing the right and left hemispheres of the WHS (generated in Edge distance calculation) were loaded into R using the package RNifti version 1.7.0 (Clayden et al., 2024). Each region was converted to a point cloud based on voxel intensity index using the R package lidR version 4.1.2 (Roussel et al., 2020). Outlying voxels (noise) were removed using the isolated voxel filter (resolution = 4, N = 25) (Roussel et al., 2020). Point clouds were converted to geometry objects in R using the package sf version 1.0.17 (Pebesma, 2018).

## Results

### Defining and describing the normative network

We calculated each individual rat cortical network as the {53×53} matrix of edge weights (*w*), representing all pairwise MIND similarity values between 53 brain regions. To estimate the normative cortical microstructural network, we computed the median of each edge weight across the 41 male adult rat brains (postnatal day [PND] 63; young adulthood) from the normative cohort (**Fig. 1A**; **Fig. 2A**; **Table 1**). The 53 regions were grouped into 15 coarse-grained cortical systems, as defined by the Waxholm Space Atlas (Kleven et al., 2023), for edge-level analyses and interpretability of results (**Fig. 1B**,**C**).

**Figure 2.**
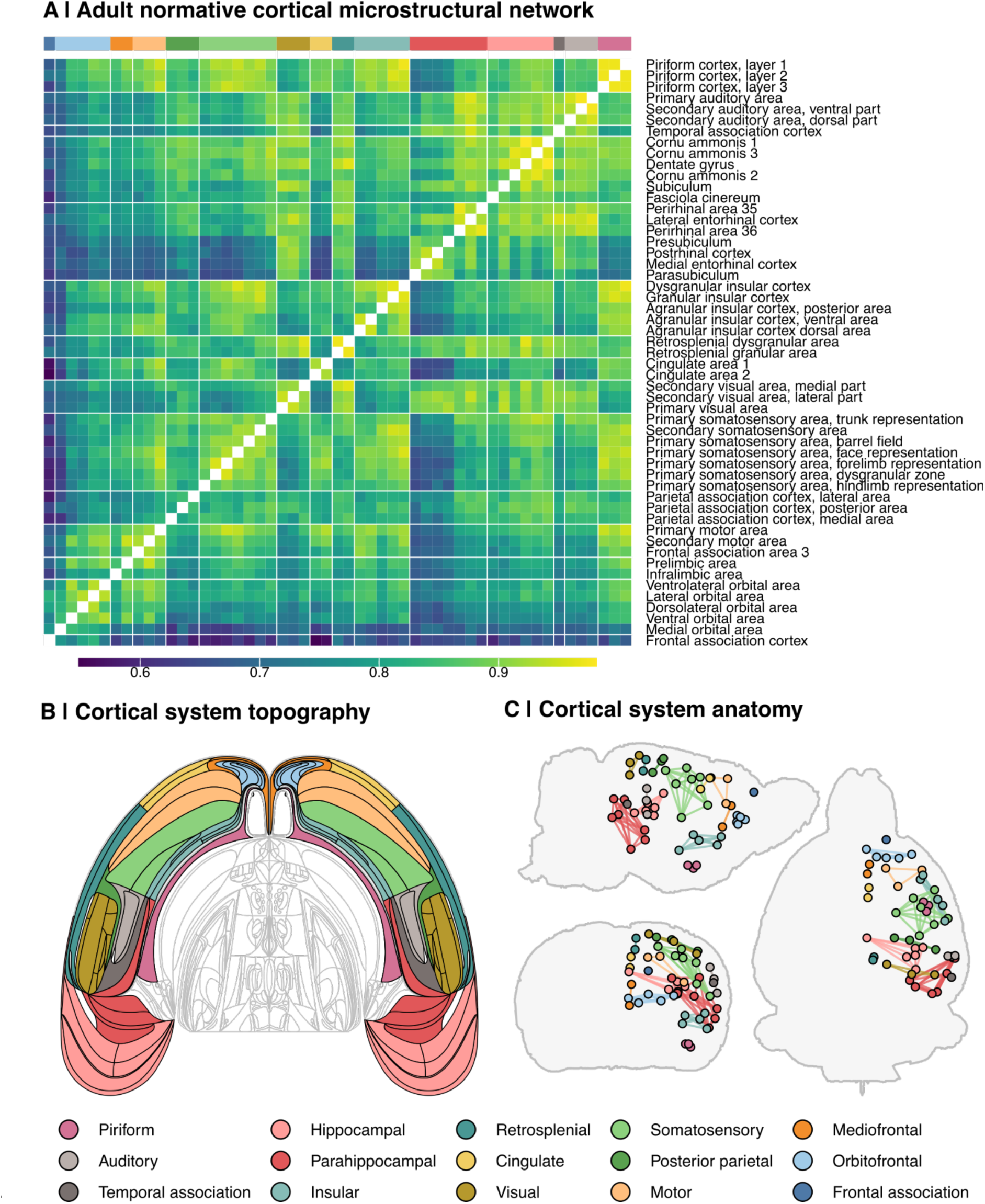
Structure of the normative rat cortical microstructural network. **A)** Heatmap representation of the normative rat connectome, defined as the median edge weight across rats in the normative cohort, at postnatal day 63 (PND 63). Rows and columns are ordered by cortical systems, as indicated by annotation bars. Tile color indicates strength of MIND similarity (edge weight, *w*). **B)** A flatmap rendering of cortical systems derived from (Hahn et al., 2021) and (Hahn and Duckworth, 2023). Colors correspond to cortical systems as indicated below. **C)** Anatomical positions of regions of interest within each cortical system. Each point represents the center for a given cortical region of interest (N=53), as defined by the WHS; lines represent intra-system edges. For coronal and axial plans, only right hemisphere nodes are shown.

### Normative network topology

The normative network distributions of nodal strength (i.e., the sum of all edge weights connecting it to the rest of the network, *s*; **Fig. 3A**) and edge weights were left-skewed (**Fig. 3B**), indicating a generally high MTR-based anatomical similarity across the cortex. Hubs, defined as regions with the highest strength (Fornito et al., 2016b), and rich-club organization, which examines whether hubs have stronger similarity with each other than expected by chance (Fornito et al., 2016a), are functionally relevant features of brain networks described in humans and other organisms (Bullmore and Sporns, 2009; Towlson et al., 2013; Rubinov et al., 2015; Swanson et al., 2024b). We found this principle held true in rat brain MIND networks. The hubs, defined as the 10 nodes with the highest strength, were largely hippocampal and piriform regions (**Fig. 3C**). The rich club coefficient (Φ), calculated using **Equation 1**, was 0.95, compared to a median coefficient of 0.45 across 10,000 distance-corrected null networks (**Fig. 3D**,**E**), indicating a significantly greater-than-random strength of connectivity amongst cortical hubs (*Z*=73; *P*_perm_=1.0*10^-4^). This evidence for a rich club organization in MIND networks was consistent with prior reports of rich clubs in the meta-analytic connectome from rat tract-tracing data (Swanson et al., 2024b) and in other species and modalities of brain networks (van den Heuvel and Sporns, 2011; Towlson et al., 2013).

**Figure 3.**
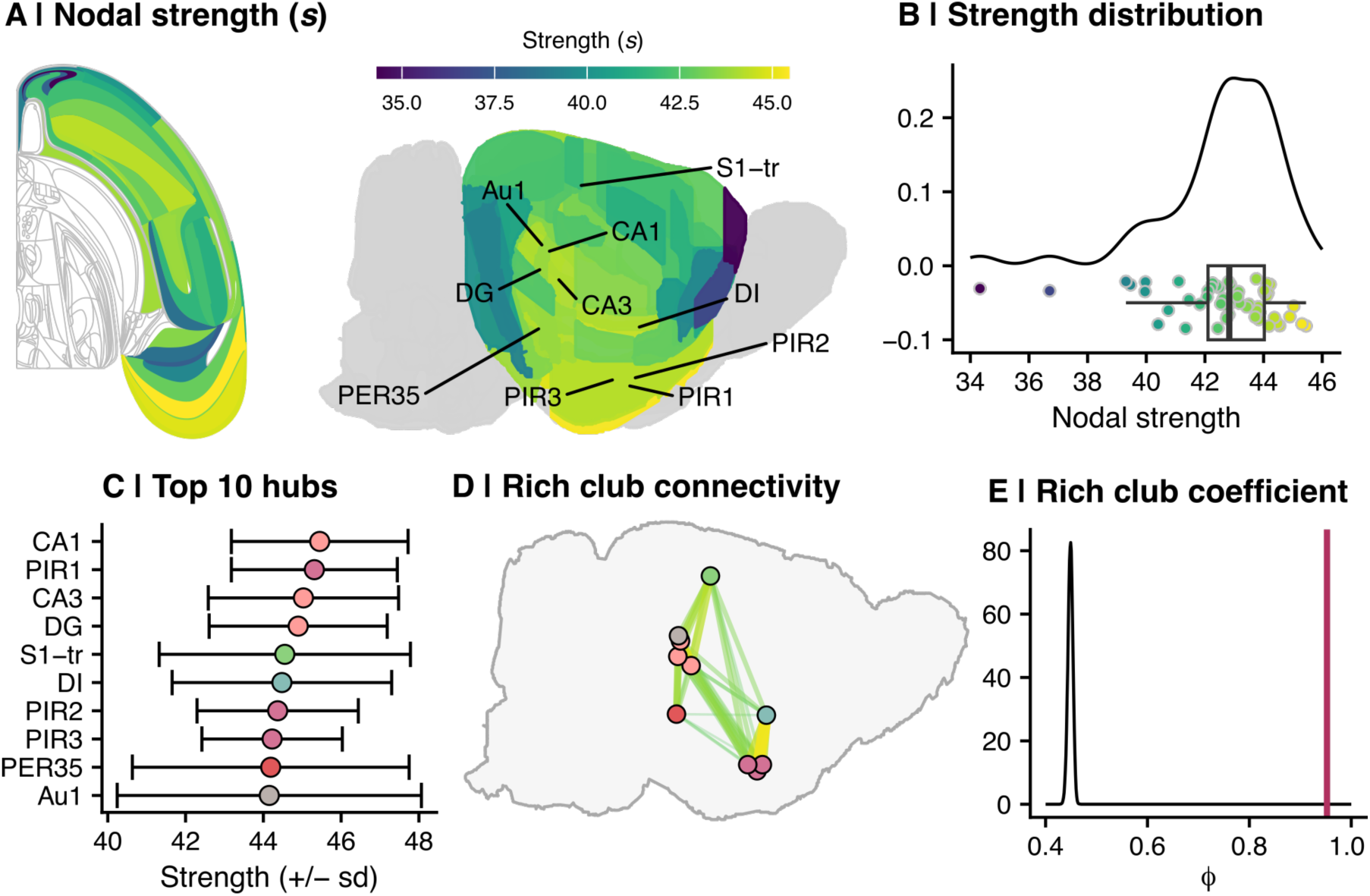
Graph-based metrics of the normative network. **A)** Maps of normative nodal strength. Left: Topographic (flatmap) cortical rendering (right hemisphere only). Right: Anatomical MRI rendering. Network hubs, or top 10 nodes by strength, are labeled. CA1=cornu ammonis 1; PIR1=piriform cortex, layer 1; CA3=cornu ammonis 3; DG=dentate gyrus; S1-tr=primary somatosensory area, trunk representation; DI=dysgranular insular cortex; PIR2=piriform cortex, layer 2; PIR3=piriform cortex, layer 3; PER35=perirhinal area 35; Au1=primary auditory area. **B)** Nodal strength distribution of the adult normative network. Each point on the x-axis shows the strength of a given region (defined as the sum of all edge weights for that region), while the y-axis shows smoothed density of nodes for each strength bin. **C)** Ten regions with the highest nodal strength, i.e., “hubs”. The x-axis represents the median strength (± standard deviation) of each region; each point is colored by cortical system as in Figure 2. The y-axis shows each region of interest (same abbreviations as panel A). **D)** Anatomical representation of the network rich club. Each point represents a hub (from panel C), and the interconnectivity among all hubs is indicated by connecting lines. The width and color of each line indicates edge weight; i.e., the strength of similiarty between the two hubs. **E)** Normative MIND network rich club coefficient versus 10000 distance-corrected null networks. The x-axis shows the rich club coefficient, while the y-axis indicates density distribution. The rich club coefficient of the normative network, defined as the sum of all hub inter-connections (N=90) divided by the sum of the top N=90 edges, is shown in by the position of the maroon line (Φ=0.95). The black curve shows the distribution of rich club coefficients across distance-corrected null networks (median Φ=0.45).

### Network validation and reliability

Research in humans has shown that structural similarity networks reflect key neurobiological features, including axonal connectivity and cytoarchitectonic similarity (Sebenius et al., 2023, 2024). Since this study is the first to describe structural similarity networks in rats, we aimed to establish these relationships within the normative network reported here. Confirming the neurobiological relevance of this network enhanced the interpretability and significance of downstream findings.

We validated the normative MIND network against four key predictions: that MIND similarity was greater between regions that (i) were spatially closer to each other, (ii) were more architectonically similar to each other, (iii) had more similar gene expression profiles, and (iv) had more similar profiles of axonal projections in prior tract-tracing data. All four of these predictions were verified as detailed below.

#### Similarity–distance relationship

First, we found that morphometric similarity decreased with increasing distance between regions such that the Euclidean distance between node centers was negatively correlated with edge weight (i.e., MIND similarity; *r*=-0.66; *P*<0.001; **Fig. 4A**). This result aligned with expectations, as brain regions that are in closer spatial proximity tend to exhibit greater structural similarity (Alexander-Bloch et al., 2013).

**Figure 4.**
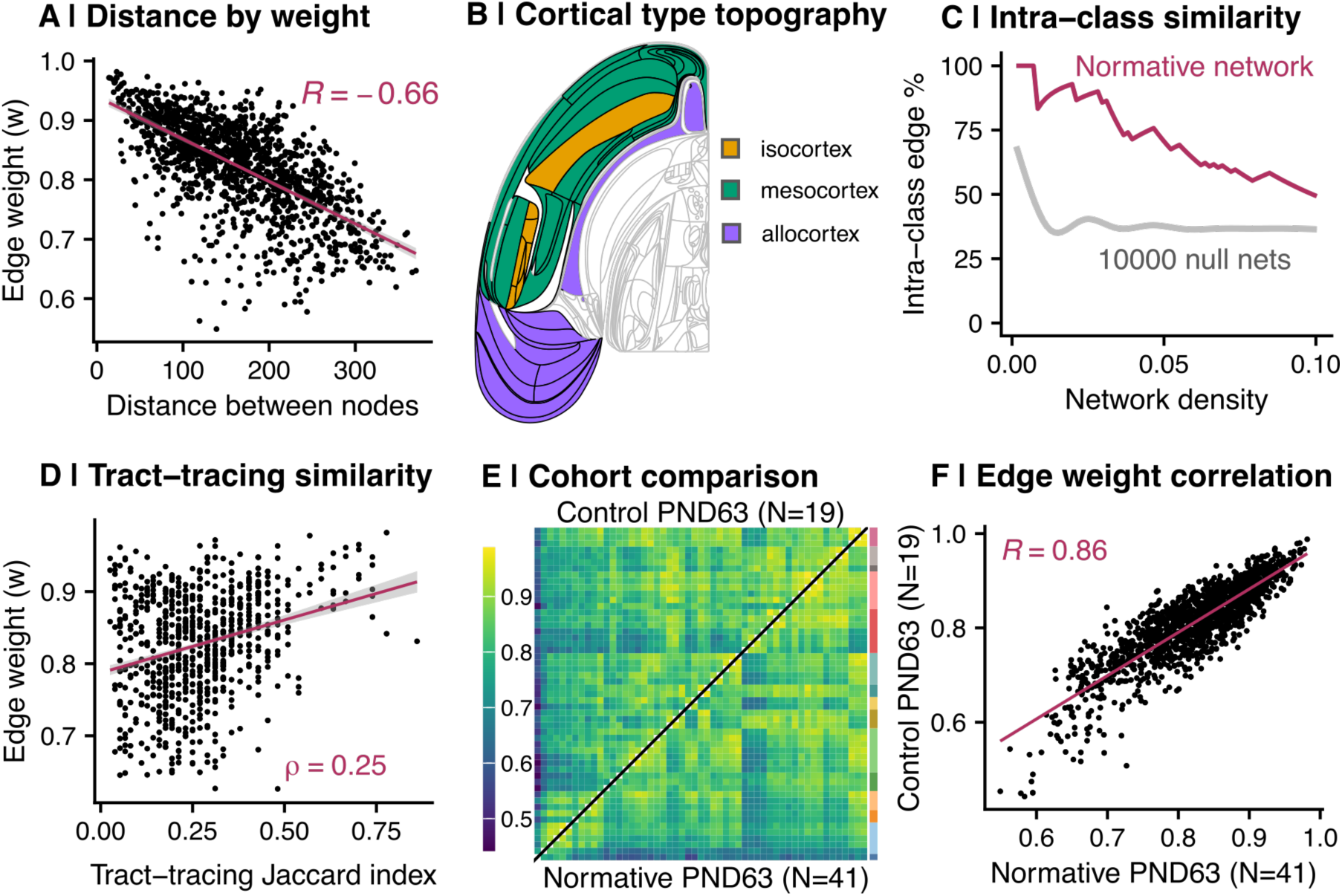
MIND-based similarity is related to metrics of cortical organization. **A)** Scatter plot of the distance between two nodes (x-axis) and their MIND similarity, or edge-weight between them (y-axis). Distance is defined as the Euclidean distance between region of interest centers, as determined using the Waxholm Space atlas. **B)** Topographic (flatmap) rendering of cortical types (left hemisphere only). Regions are colored by cortex type, as defined by (García-Cabezas et al., 2023). **C)** Proportion of intra-class edges across network density thresholds. The x-axis shows the proportion of top-weighted edges considered, while the y-axis indicates the percentage of those top-weighted edges that consists of two regions from the same cortex class. The maroon curve shows the percentage of within-class edges in the normative MIND network, while the gray line shows the mean percentage (± standard deviation) of within-class edges across 10000 permuted null networks. **D)** Correlation between similarity of tract-tracing connection profiles between pairwise combinations of regions and strength of MIND similarity. The Jaccard index for edge *ij* was defined as the intersection of tract-tracing connections (i.e., where *ik* == *jk*) divided by the union of all connections containing *i*, *j*, or both (**Equation 2**). **E)** Heatmap representations of the median PND 63 control network from the experimental stress cohort (top left of the diagonal) and the median PND 63 network from the normative developmental cohort (bottom right of the diagonal). Panel legend is the same as Figure 2A. **F)** The relationship between edge weights in the median PND 63 network from the normative development cohort (x-axis) and median PND 63 control network from the experimental stress cohort (y-axis; *r*=0.86; *P*<0.001). Each point represents an edge; the line of best fit is shown in maroon.

#### Cytoarchitectonic relationships to similarity

Second, if MIND similarity truly reflects cytoarchitectonic similarity, we would expect regions of the same cortical type to exhibit higher MIND edge weights. The rat neocortex is structured along gradients of laminar differentiation, which expanded from allocortical areas (archicortical [hippocampal] and paleocortical [olfactory]) according to the dual origin theory (García-Cabezas et al., 2023) (**Fig. 4B**). In rodents, these cortical gradients extend through mesocortical (agranular and dysgranular) into isocortical (eulaminate) areas (García-Cabezas et al., 2023). To test the relationship between MIND similarity and cytoarchitectonics, we co-registered our neuroimaging data with a cytoarchitectonic atlas of the rat brain (García-Cabezas et al., 2023) (**Fig. 4B**), and compared MIND edge weights between areas of the same cortex class (“intra-class” edges) versus areas of different cortex classes (“inter-class” edges), removing isocortical regions due to small sample size (N=5 regions).

Intra-class edges between two allocortical areas (“allo-allo”) had significantly higher MIND similarity than edges between allocortical and mesocortical (“allo-meso”) areas (false discovery rate, FDR < 0.05; (**Fig. S2**). However, meso intra-class edges were not significantly more similar than allo-meso. To probe this relationship further, we calculated the percentage of intra-class edges (allo-allo and meso-meso) at various binary network densities, thresholding across a range of sparse connection densities (1% to 10% of all possible pairwise edges). We then compared these to distance-corrected null networks thresholded across the same range (**Fig. 4C**). Across all thresholds, the empirical rat network demonstrated a higher percentage of intra-class edges than the null networks (**Fig. 4C**), indicating a higher-than-chance density of cytoarchitectonically similar edges among edges with the highest MIND similarity.

#### Mouse transcriptional relationships to similarity

Third, we predicted that regions with more similar patterns of gene expression would also exhibit higher MIND similarity. However, a spatial transcriptomic atlas of the rat brain is not yet available. To overcome this limitation, we leveraged homology between the rat and mouse brain to access spatially comprehensive transcriptomic data from the Allen Mouse Brain Atlas (AMBA; N=437 genes) (Yao et al., 2023). While we acknowledge the challenges of cross-species comparisons, our goal was to assess the general consistency between edge-wise MIND similarity and transcriptional similarity, rather than expecting a precise one-to-one correspondence.

We tested and verified that regions with higher MIND similarity also demonstrated higher transcriptomic similarity, calculated as the pairwise regional correlation between gene expression profiles (**Fig. S3**). These findings support our general hypothesis that MIND similarity reflects broad patterns of transcriptomic similarity.

#### Tract-tracing relationships to similarity

Fourth, we hypothesized that MIND similarity would reflect patterns of axonal connectivity. To test this, we converted meta-analytic rat tract-tracing data (Swanson et al., 2024b) to a similarity matrix by calculating the Jaccard index of the log10-transformed ordinal connection weight between pairwise regions (**Equation 2**; **Fig. S4A**). This transformation yielded a continuous measure representing the similarity of axonal connectivity profiles between regions (i.e., the extent to which two regions share common connectivity patterns).

When the tract-tracing Jaccard index was correlated with MIND edge weights, we found a positive relationship (ρ=0.25; *P*<0.001; **Fig. 4D**). To assess the significance of this correlation, we compared it to a null distribution generated from 10000 distance-corrected null networks. The observed Spearman correlation was significantly greater than chance (*Z*=6.04; *P*_perm_=1.0*10^-4^; **Fig. S4B**), suggesting a meaningful relationship between anatomical connectivity profiles and MIND similarity.

#### Cross-cohort reproducibility

Finally, to assess the reliability of MIND network analysis, we directly compared the median adult MIND network for the normative developmental cohort (N=41, PND 63; **Fig. 1A**) to the median adult MIND network for an independent cohort of rats: namely, the normally reared (control) group in the experimental stress cohort (N=19, PND 63; **Fig. 1B**; **Fig. 4E**). Edge weights were highly correlated between the normative adult and independent control MIND networks (*r*=0.86; *P*<0.001; **Fig. 4F**), as was nodal strength (*r*=0.91; *P*<0.001). These results demonstrate a high level of replicability of the cortical microstructural network estimated by identically implemented MIND analyses of MTR data collected using the same sequences in two independent cohorts of rats.

### Normative developmental changes in the rat cortical microstructural network

Having established convergent validity and inter-sample replicability of MIND networks, we next harnessed this analytic approach to model network level reorganization of the rat brain over development. This strategy, applied to a longitudinal MRI dataset spanning PND 20, 35, 63 and 230 (N=162 total scans, **Fig. 1A**, **Methods**, **Table 1**), revealed developmental changes in (i) each region’s morphometric similarity with the rest of the brain (strength, *s*), and (ii) the morphometric similarity (edge strength, *w*) between each unique pair of regions. Visual inspection of the median MIND matrices for each of the four time-points indicated that there are age-related changes in the cortical pattern of inter-areal similarity. For example, frontal cortical areas (frontal association cortex, orbitofrontal cortex, mediofrontal cortex, and motor areas) become more similar to the rest of the cortex during early development, and then strikingly less similar during later aging (**Fig. 5A**). Likewise, nodal strength showed a general tendency to increase during development and decrease during aging (**Fig. 5B**).

**Figure 5.**
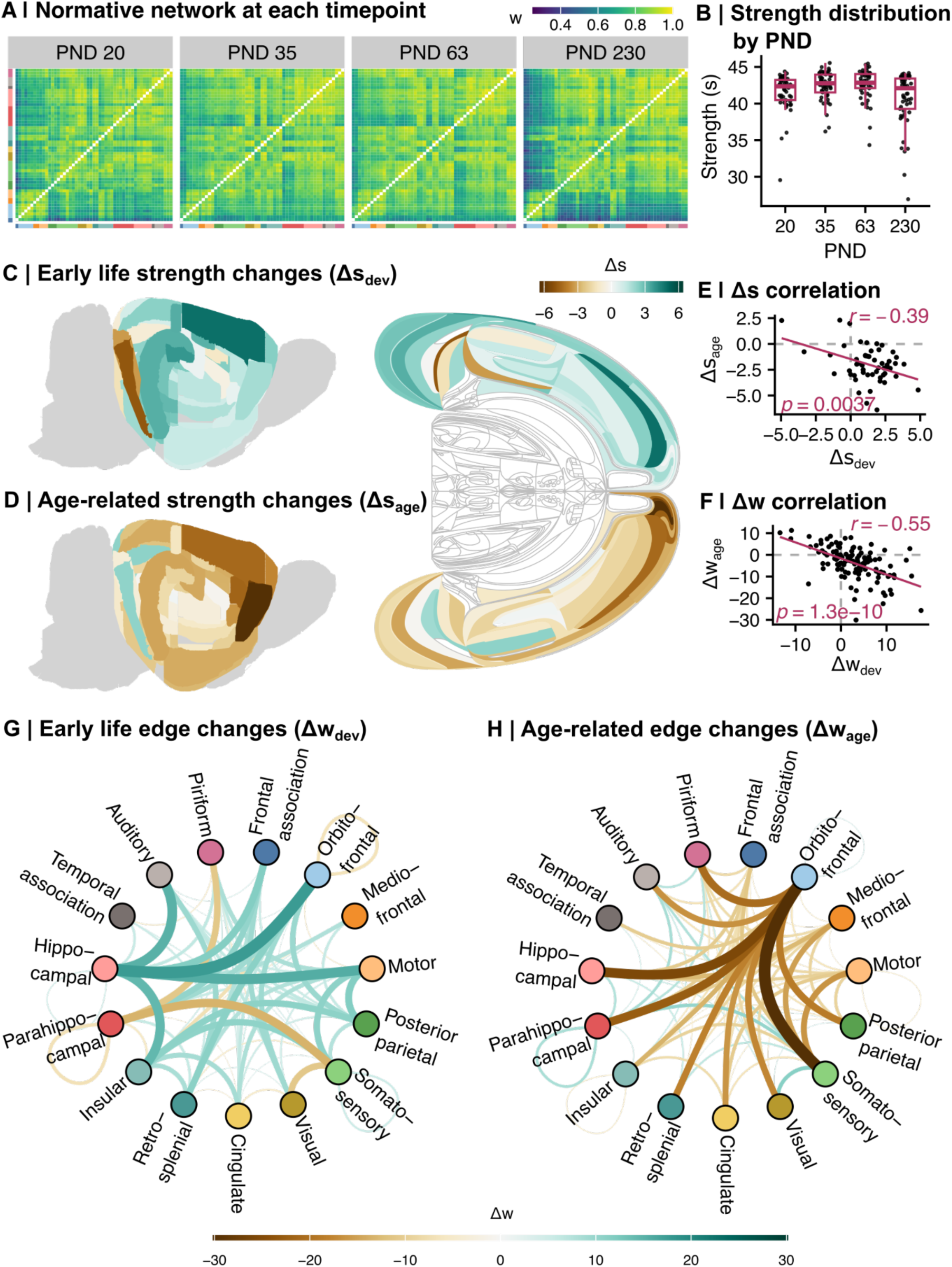
The normative rat cortical connectome generally increases in similarity in early development and decreases in similarity in aging. **A)** Heatmap representation of the connectome throughout development in the normative cohort (median edge weight, at each timepoint). Rows and columns are ordered by decreasing-to-increasing nodal strength within broader cortical system (in the same order as **Figure 2A**). Tile color indicates strength of MIND similarity (edge weight, *w*). **B)** The normative strength distribution through development, defined as the median strength per ROI at each timepoint. Each point represents a region of interest; boxplots show the overall strength (*s*) distribution at each timepoint. The normative strength distribution at PND 20 is significantly lower than the distribution at PND 63 (Dunn test *P*=0.019); the nodal strength distribution at PND 230 is significantly lower than the distributions at PND 35 (Dunn test *P*=0.027) and PND 63 (Dunn test *P*=0.003) (Krustal-Wallis *P*=0.002). **C)** Anatomical patterning of nodal strength changes in early development (Δs_dev_). Left: Volumetric rendering; Right: Flatmap rendering (left hemisphere/top half only; (Hahn et al., 2021; Hahn and Duckworth, 2023)). Brown indicates decrease in strength, teal indicates increase in strength. Δs_dev_ was considered significant if |*t*-value| of the age term in the mixed effects model was greater than 2. **D)** Anatomical patterning of nodal strength changes in aging (Δs_age_; flatmap rendering right hemisphere/bottom half). Figure key same as panel C. **E)** Pearson correlation between Δs_dev_ (x-axis) and Δs_age_ (y-axis; *r*=-0.39; *P*=0.004). Each point represents a region of interest; the line of best fit is shown in maroon. **F)** Pearson correlation between Δw_dev_ (x-axis) and Δw_age_ (y-axis; *r*=-0.55; *P*<0.001). Each point represents an edge; the line of best fit is shown in maroon. **G)** A circle plot of significantly changed system-level edges in early development (Δw_dev_; |*t*|> 3.3). Each datapoint in the circle represents a broader cortical system, colored and labeled by system. The color and size of the curves connecting two points show the change in edge weight (brown indicates decreasing similarity; teal indicates increasing similarity). **H)** A circle plot of significantly changed edges in aging (Δw_age_). Figure key same as panel G.

We quantified the regional rate of change in early development as the linear gradient or slope of age-related change in nodal strength for each cortical area between PND 20 (weanling) to PND 35 (adolescence), Δs_dev_, accounting for inter-subject variation (**Equation 3A**). Similarly, to calculate changes in aging, we applied the same model to estimate the age-related change in strength for each cortical area from PND 63 (young adult) to PND 230 (mid adult) timepoints, Δs_age_.

During development, regions in most cortical systems significantly increased in nodal strength, especially the hippocampal formation and motor cortex (Δs_dev_ *t*-values=7.19 and 5.22, respectively; **Fig. 5C**). The parahippocampal region decreased in similarity with the rest of the brain (Δs_dev_ *t*=-1.34). In contrast to these developmental changes, most cortical systems decreased in nodal strength during aging, especially areas of the frontal cortex, including the orbitofrontal (Δs_age_ *t*=-11.4), motor (Δs_age_ *t*=-6.13), and mediofrontal (Δs_age_ *t=*-5.94; **Fig. 5D**) cortices. Early life and aging effects on nodal strength were negatively correlated (*r*=-0.39; *P*=0.004; **Fig. 5E**), indicating that those regions showing the strongest strength increases in development also tended to show the most rapid strength decreases in aging. Ten fronto-hippocampal regions (Frontal: dorsolateral orbital area, frontal association cortex, frontal association area 3, primary motor area, secondary motor area; Hippocampal: dentate gyrus, perirhinal area 35, cornu ammonis 1) both significantly increased in strength in early development and significantly decreased in strength in aging. Only one region, the parasubiculum, significantly decreased in early development and increased in aging.

We also calculated changes in early development and aging for each inter-areal similarity or edge in the MIND networks (Δw_dev_ and Δw_age_, respectively; **Equation 3B**). In early development, the hippocampal formation demonstrated notable increases in similarity with frontal regions, including the orbitofrontal (Δw_dev_ *t=*17.6), motor (Δw_dev_ *t*=15.5), frontal association (Δw_dev_ *t=*11.7), and mediofrontal (Δw_dev_ *t*=7.97) cortices (**Fig. 5G**). In contrast, parahippocampal regions largely decreased in similarity with other cortical systems, including the somatosensory (Δw_dev_ *t*=-13.4) and piriform (Δw_dev_ *t*=-11.0) cortices. In aging, the most salient changes in edge weight involved orbitofrontal regions, which significantly diverged from every other cortical system, most notably the somatosensory (Δw_age_ *t*=-30.1), hippocampal (Δw_age_ *t*=25.7), parahippocampal (Δw_age_ *t*=22.5) regions (**Fig. 5H**). As with strength, edge weight effects during development and aging were negatively correlated (*r*=-0.55; *P*<0.001; **Fig. 5F**), demonstrating that frontal and hippocampal systems with the most rapid increases in pairwise similarity during early development showed the fastest decreases in similarity, or increases in dissimilarity, during aging.

### Early life environmental stressors perturb adult cortical similarity networks

We assessed the impact of early life stress—as modeled by repeated maternal separation (RMS)—on the nodal strength of the young adult cortical microstructural network by running multiple case-control comparisons for each cortical area at the PND 63 timepoint (**Equation 4A**; **Fig. 1B**). Exposure to RMS was associated with strength decreases in most brain regions (N=40 of 53 total) (**Fig. 6A**). Four regions had significant differences following permutation testing: the lateral entorhinal cortex, perirhinal area 36, and cingulate area 1 decreased in strength in RMS (permutation *Z*-scores=-3.07, -2.31, and -2.18, respectively), while the frontal association cortex demonstrated increased strength following RMS (*Z*_perm_=2.08; **Fig. 6B**)

**Figure 6.**
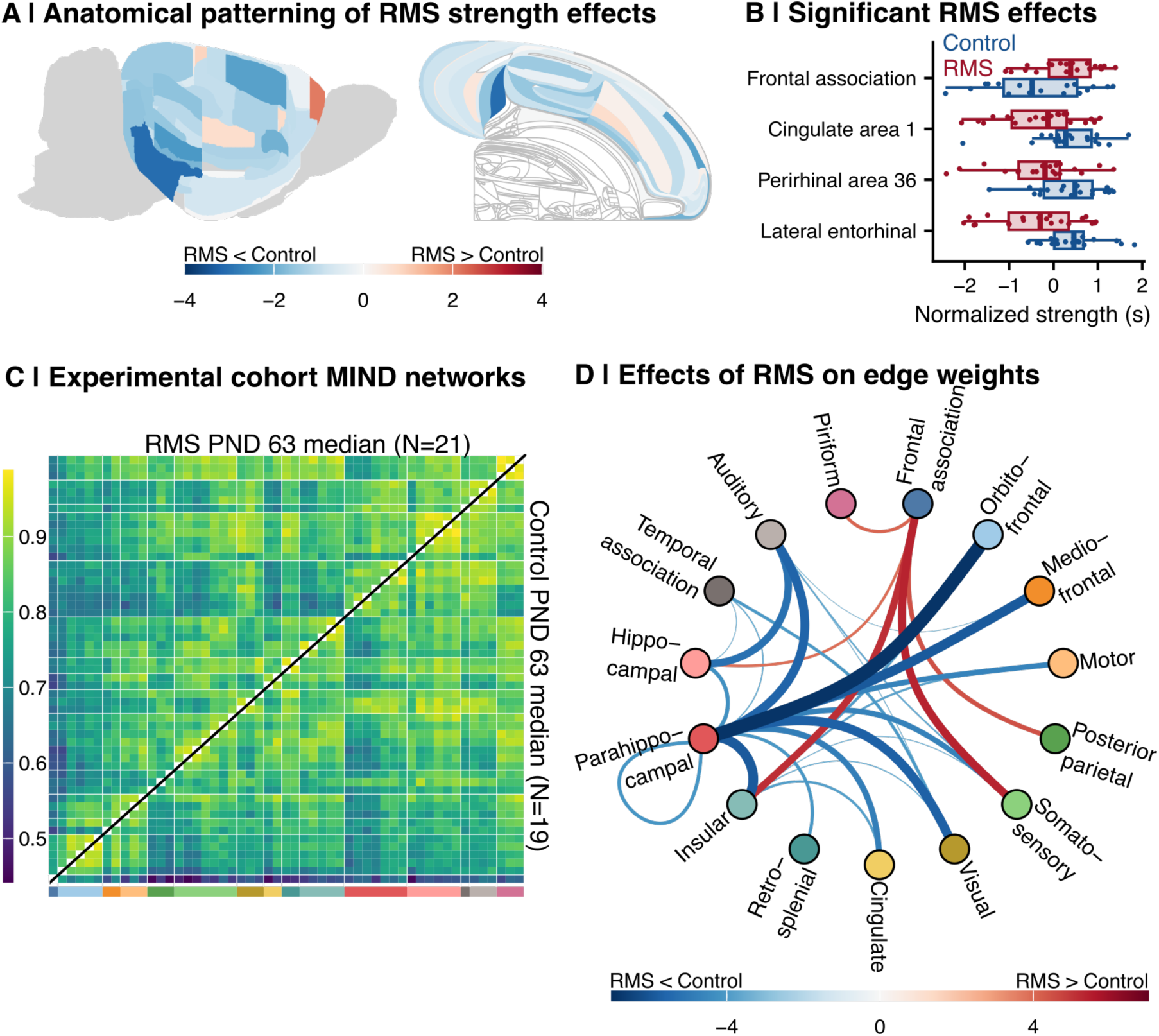
Impact of early life stress on nodal strength and edge weights. **A)** Anatomical distribution of the post-natal day 63 (young adulthood) RMS-control effect sizes across brain slices. For each region of interest, a linear model was run of the normalized strength on group + age + sex + TBV, and the group statistic was extracted as the actual effect size. Then, 1000 permutations were run in which the group assignments were shuffled, the same linear model was run, and the group statistic was extracted as the permuted effect size. The *Z*-score of the actual effect size was calculated as its position in the permuted distribution. Regions with positive *Z*-scores (shown in red) demonstrated increased strength in RMS subjects in young adulthood; regions with negative *Z*-scores (shown in blue) demonstrated decreased strength. The flatmap rendering was derived from (Hahn et al., 2021; Hahn and Duckworth, 2023). **B)** Boxplots showing the nodal strength distribution by group for the four regions with significant case-control differences at PND 63 (|*Z*_perm_| > 1.96). The x-axis shows nodal strength, corrected for covariates and normalized for visualization purposes. Each point represents a subject, while the box-and-whiskers plot shows the overall distribution by group. Blue indicates control, red indicates RMS. **C)** The median PND 63 MIND networks in control subjects (top) vs RMS subjects (bottom). Figure legend is the same as **Fig. 2A. D)** A circle plot of significant RMS-control edge weight differences in young adulthood (PND 63; |*t*| > 3.3). Node order and system annotation are the same as Figure 5 panels G and H. Curves connecting two points indicate a significant case-control difference in edge weight; colored by effect size (with red indicating higher weight in RMS and blue indicating higher weight in control); line width indicates absolute value effect size.

Qualitatively, visual inspection of the median MIND matrices in control (N=19) and RMS-exposed rats (N=21) indicated that the early life stressor induced changes in the cortical pattern of inter-areal similarity, especially in frontal and hippocampal systems (**Fig. 6C**). Indeed, edge-level case-control analyses (**Equation 4B**) showed that RMS-exposed rats had strongly decreased similarity between parahippocampal cortex and several other cortical areas (including the orbitofrontal cortex (*t=*-6.89), mediofrontal cortex (*t=*-5.89), and motor cortex (*t*=-4.69; **Fig. 6D**). This contrasts with markedly increased similarity between the frontal association cortex and other areas, including the somatosensory (*t*=5.40) and insular cortices (*t*=5.17). Together, these results indicate vulnerability in network structure to environmental stress, especially in frontal and parahippocampal regions most sensitive to developmental and aging changes in nodal strength.

### Effects of early life stress are nested within normative developmental changes in the cortical microstructural network

Finally, we tested more formally for potential congruence between stress effects and normative network changes in development and aging. Strikingly, variation in the effects of RMS on inter-areal edge weights in the rat cortical microstructural network was positively correlated with variation in the edge weight changes over normative development (Δw_dev_; *r*=0.18, *P*_perm_=0.03; *Z*_perm_=1.90; **Fig. S5** (left)). Specifically, those edges showing greatest similarity increases in normative development also tended to show greater similarity increases following RMS. Furthermore, the effects of RMS on edge weight were significantly and negatively correlated with normative age-related changes in edge weight (Δw_dev_; *r*=-0.19, *P*_perm_=0.02; *Z*_perm_=-2.03; **Fig. S5** (right)), indicating that the edges with the greatest similarity increases following RMS were also those that decreased in similarity the most in normative aging. Taken together, these results are consistent with a model in which early life stress accelerates normative brain reorganization during development; and areas which are most developmentally dynamic and vulnerable to stress are also the most susceptible in aging.

## Discussion

We have pioneered and validated MIND similarity network analysis as a novel approach to inferring myelo-architectonic similarity between all cortical areas in an individual rat’s brain. Using this methodological advance, we studied N=47 rat cortical networks to investigate normative dynamics in cortical similarity across development and aging. These changes, which primarily involved frontal and hippocampal systems, were examined through MIND analysis of myelin-sensitive MTR data collected up to four times over each rat’s lifespan. In a second experiment, we tested the hypothesis that early life stress exposure was associated with subsequent abnormalities of cortical network organization. Repeated maternal separation in the first 20 days of postnatal life affected the adult similarity of cortical areas, especially frontal and hippocampal systems that are normatively most dynamic in adolescent development and early brain aging.

Structural MRI similarity analysis is being increasingly used as a measure of brain network organization in various methodological and experimental contexts ((Sebenius et al., 2024) and references therein). In general, the interpretation of MIND similarity rests on two key assumptions: (i) that the correlation or inverse divergence between MRI features in two cortical areas reflects their cyto-architectonic or myelo-architectonic similarity; and (ii) that cortical areas which are more architectonically similar are more likely to be axonally inter-connected (Bazinet et al., 2023; Sebenius et al., 2024). We can therefore think of a structural MRI similarity network as primarily a map of cortical patterning—an architectome—which is in turn a partial proxy for the map of axonal wiring—a connectome. To validate rat brain MIND similarity analysis, we tested both these key assumptions. The results confirmed that cortical areas belonging to the same architectonic class, or spatially adjacent to each other, tended to have higher MIND similarity, and that higher MIND similarity between cortical areas was associated with stronger evidence of similarity in axonal connectivity, based on a prior meta-analysis of tract-tracing studies (Swanson et al., 2024b). Since the MRI data we used for this analysis were collected using a magnetization transfer (MTR) sequence that is known to be sensitive to cortical myelination and neuropil density (Grossman et al., 1994), we can therefore generally interpret MTR-derived MIND (dis-)similarity as indicative of architectonic differentiation and myelination of the cortex of an individual rat.

To characterize the dynamics of such differentiation across the lifespan, we measured changes in similarity in early life, defined as PND 20 to 35, and in later life, defined as PND 63 to 230. Though the data on developmental patterns of rat brain myelination are sparse, a histological study demonstrated that the rat brain begins myelinating around PND 10, and most areas are fully myelinated by PND 24 (Downes and Mullins, 2014). Thus, our “early” developmental data likely reflect the end of early life myelinating processes (e.g., late adolescence) and do not capture earlier peaks in cortical myelination. We show that increasing MIND similarity in fronto-hippocampal circuitry in late adolescent development (“early life”) is coupled to rapidly decreasing similarity of these systems in mid-adulthood aging. These data thus support the “last in, first out” hypothesis of development and aging, in which plastic regions that mature last in early development (namely, frontal areas related to decision-making and executive function) are more vulnerable to age-related decline, as previously described in humans (Douaud et al., 2014; Fjell et al., 2014; Duan et al., 2024). We demonstrate here that rat brain microarchitecture is subject to the same phenomenon.

Developmentally sensitive fronto-(para)hippocampal circuitry also showed targeted disruptions in young adult rats who had been exposed to repeated maternal separation. This result aligns with studies in humans, in which later-developing plastic circuitry also shows increased vulnerability to disease processes during early development and aging, such as schizophrenia and Alzheimer’s disease, respectively (Douaud et al., 2014; Fjell et al., 2014). The congruence between developmentally dynamic and environmentally sensitive regions was reinforced quantitatively, as edges that increased in similarity in development tended to show higher similarity following RMS (**Fig. S5**). These results are consistent with an accelerated development hypothesis of early life stress (Callaghan and Tottenham, 2016). Edges that showed higher similarity in young adulthood following early life stress also demonstrated more rapid divergence in normative aging. This provides further support for the concept that specific fronto-hippocampal circuits are developmentally dynamic, vulnerable to age-related decline, and susceptible to environmental stress (Douaud et al., 2014; Duan et al., 2024). The alignment between human and rat data underscores the potential of this model system for experimental investigation into the mechanisms linking early life stress, network development, and aging.

An increase in similarity between two areas (in this case, frontal and hippocampal) does not necessarily indicate the formation of new axonal connections but likely represents coordinated changes in (i) myelination of fibers, and/or (ii) microstructural properties, such as synaptic or dendritic remodeling. The first hypothesis is supported by rat histological data showing that, between PND 24 and PND 37, myelination occurs exclusively in the fornix and mammillothalamic tract—pathways that traverse the hippocampus (Downes and Mullins, 2014). It has been argued theoretically that regions related to memory and learning develop as the adolescent rat ventures out and requires spatial recognition, a skill less essential earlier in life (Downes and Mullins, 2014). Consistent with this, a study on rat hippocampal myelination reported the first appearance of myelinated fibers at PND 17, a significant increase to near-adult levels by PND 25, and full maturation by PND 60 (Meier et al., 2004).

In addition to myelination, changes in MIND similarity may also arise from coordinated microstructural reorganization. Extensive cross-modality research highlights hippocampal plasticity in rodents, especially changes in synaptic density in response to environmental enrichment or deprivation (Lerch et al., 2011; Stein et al., 2016; Ohline and Abraham, 2019; Bramati et al., 2023; Dayananda et al., 2023). Similarly, the frontal cortex undergoes environmentally-sensitive synaptogenesis in early development—continuing into adulthood in rats (Zeiss, 2021)—and synaptic pruning throughout the lifespan (Kolk and Rakic, 2022). These shifting neuropil profiles suggest that cortical regions without direct physical connections may exhibit high MIND similarity due to convergent synaptic architectures—or increasingly divergent profiles with other regions. Our cross-validation data support this concept. In the tract-tracing (Swanson et al., 2024b) Jaccard comparison, a subset of edges showed low tract-tracing similarity but high MIND similarity, predominantly involving hippocampal regions. Excluding hippocampal edges strengthened the correlation between MIND similarity and tract-tracing (ρ=0.41; *P*<0.001; **Fig. S4C**). We postulate that the observed high MIND similarity, despite low axonal connection similarity, may reflect convergent plastic reorganization of the microstructural properties of these regions. Notably, the hypotheses of cortical fiber myelination and synaptic density as contributors to MIND similarity are likely interconnected, as evidence suggests neural activity can induce myelinogenesis (Demerens et al., 1996). Future studies incorporating histological measures of synaptic density and activity-dependent myelination in the rat brain could help clarify the relative contributions of these mechanisms to MIND similarity.

A key limitation of this study is that the normative developmental cohort included only male animals; accordingly, the normative network defined here represents male development only. To more fully characterize normative development, future work should explicitly investigate sex differences in network structure and dynamics. It would also be of interest to reapply these methods using varying parcellation schemes, as different definitions of brain regions—both cortical and subcortical—may offer complementary or more detailed insights. Future work to characterize MRI-derived rat brain network architecture could include additional morphometric features, such as DTI metrics, within a multivariate MIND analysis. This may provide a broader view of cortex-wide morphological co-variation and more directly reflect axonal connections. As additional rat brain resources become available (for instance, a brain-wide spatial transcriptomic atlas and consensus nomenclature/parcellations), our ability to biologically annotate the rat structural similarity network will continue to improve.

We provide the normative MTR-derived rat cortical microstructural network and all associated code to support further investigation and understanding of the complex organization of cortical networks in this key model system. Our results demonstrate the biological validity and replicability of the MIND similarity analysis and demonstrate its sensitivity to developmental and environmental stress-related changes in cortical network configuration. We emphasize the importance and vulnerability of key frontal and hippocampal circuitry in dynamic processes of normative development and atypical developmental trajectories triggered by early life adversity.

## Data and Code Availability

All data were made available through previous publications (Dutcher et al., 2023; Jones et al., 2024). All visualizations were rendered using the R package ggplot version 3.5.1 (Wickham, 2016). Code for data preprocessing (including the rat MRI preprocessing pipeline), data analysis, and figure construction is available at https://github.com/rlsmith1/rat_MRI_similarity_networks.

## Author Contributions

*Designed research*: R.L.S., E.G.D., J.A.J., J.W.D., F.J.M., A.R., P.E.V., E.T.B., *Performed research*: R.L.S., S.J.S., L.D., E.G.D., J.A.J., *Contributed unpublished reagents/analytic tools*: S.J.S., J.D.H., O.S., L.W.S., P.A.T., D.R.G., *Analyzed data*: R.L.S., *Wrote the paper*: R.L.S., S.J.S., L.D., E.G.D., J.A.J., J.D.H., O.S., L.W.S., P.A.T., D.R.G., J.W.D., F.J.M., A.R., P.E.V., E.T.B.

## Acknowledgments

R.L.S completed this work as a PhD candidate in the NIH Oxford-Cambridge Scholars Program. R.L.S., P.A.T., D.R.G., A.R., and F.J.M NIMH Intramural Research Program of the NIH/HHS, USA as follows: R.L.S and F.J.M (1ZIAMH002810), A.R. (1ZIAMH002949), P.A.T. and D.R.G (ZICMH002888). L.D. and E.G.D. were supported by the Gates Cambridge Scholarship. P.E.V. was supported by MQ: Transforming Mental Health (MQF17_24). This work received computational support from the NIP HPC Biowulf cluster (http://hpc.nih.gov). All research from the Department of Psychiatry at the University of Cambridge is made possible by the mental health theme of the NIHR Cambridge Biomedical Research Centre (BRC-1215-20014) and the NIHR East of England Applied Research Centre. E.T.B. was also supported by an NIHR Senior Investigator award. The views expressed are those of the author(s) and not necessarily those of the NHS, the NIHR or the Department of Health.

## References

Abraham M, Peterburs J, Mundorf A (2023) Oligodendrocytes matter: a review of animal studies on early adversity. J Neural Transm (Vienna) 130:1177–1185.

Alexander-Bloch A, Giedd JN, Bullmore E (2013) Imaging structural co-variance between human brain regions. Nat Rev Neurosci 14:322–336.

Bai LS, Kinosada Y, Okuda Y, Ning M, Nakagawa T (1996) Changes of magnetization transfer ratio according to rat brain development. Nihon Igaku Hoshasen Gakkai Zasshi 56:955– 960.

Bass NH, Netsky MG, Young E (1970) Effect of neonatal malnutrition on developing cerebrum. II. Microchemical and histologic study of myelin formation in the rat. Arch Neurol 23:303– 313.

Bazinet V, Hansen JY, Vos de Wael R, Bernhardt BC, van den Heuvel MP, Misic B (2023) Assortative mixing in micro-architecturally annotated brain connectomes. Nat Commun 14:2850.

Benjamini Y, Hochberg Y (1995) Controlling the false discovery rate: a practical and powerful approach to multiple testing. Journal of the royal statistical society series b-methodological 57:289–300.

Bramati G, Stauffer P, Nigri M, Wolfer DP, Amrein I (2023) Environmental enrichment improves hippocampus-dependent spatial learning in female C57BL/6 mice in novel IntelliCage sweet reward-based behavioral tests. Front Behav Neurosci 17:1256744.

Breton JM, Barraza M, Hu KY, Frias SJ, Long KLP, Kaufer D (2021) Juvenile exposure to acute traumatic stress leads to long-lasting alterations in grey matter myelination in adult female but not male rats. Neurobiol Stress 14:100319.

Bryda EC (2013) The Mighty Mouse: the impact of rodents on advances in biomedical research. Mo Med 110:207–211.

Bullmore E, Sporns O (2009) Complex brain networks: graph theoretical analysis of structural and functional systems. Nat Rev Neurosci 10:186–198.

Bullmore E, Sporns O (2012) The economy of brain network organization. Nat Rev Neurosci 13:336–349.

Callaghan BL, Tottenham N (2016) The Stress Acceleration Hypothesis: effects of early-life adversity on emotion circuits and behavior. Current Opinion in Behavioral Sciences 7:76–81.

Cao H, Wei P, Huang Y, Wang N, Guo L-A, Fan X, Wang Z, Ren L, Piao Y, Lu J, Shan Y, He X, Zhao G (2023) The alteration of cortical microstructure similarity in drug-resistant epilepsy correlated with mTOR pathway genes. EBioMedicine 97:104847.

Clayden J, Cox B, Jenkinson M, Hall M, Reynolds R, Fissell K, Gailly J-L, Adler M (2024) RNifti: Fast R and C++ Access to NIfTI Images. Available at: https://cran.r-project.org/web/packages/RNifti/index.html [Accessed January 30, 2025].

Cox RW (1996) AFNI: Software for Analysis and Visualization of Functional Magnetic Resonance Neuroimages. Comput Biomed Res 29:162–173.

Dayananda KK, Ahmed S, Wang D, Polis B, Islam R, Kaffman A (2023) Early life stress impairs synaptic pruning in the developing hippocampus. Brain Behav Immun 107:16–31.

Demerens C, Stankoff B, Logak M, Anglade P, Allinquant B, Couraud F, Zalc B, Lubetzki C (1996) Induction of myelination in the central nervous system by electrical activity. Proc Natl Acad Sci U S A 93:9887–9892.

Dorfschmidt L, Váša F, White SR, Romero-García R, Kitzbichler MG, Alexander-Bloch A, Cieslak M, Mehta K, Satterthwaite TD, NSPN Consortium, Bethlehem RAI, Seidlitz J, Vértes PE, Bullmore ET (2024) Human adolescent brain similarity development is different for paralimbic versus neocortical zones. Proc Natl Acad Sci U S A 121:e2314074121.

Douaud G, Groves AR, Tamnes CK, Westlye LT, Duff EP, Engvig A, Walhovd KB, James A, Gass A, Monsch AU, Matthews PM, Fjell AM, Smith SM, Johansen-Berg H (2014) A common brain network links development, aging, and vulnerability to disease. Proc Natl Acad Sci U S A 111:17648–17653.

Downes N, Mullins P (2014) The development of myelin in the brain of the juvenile rat. Toxicol Pathol 42:913–922.

Duan H et al. (2024) Population clustering of structural brain aging and its association with brain development. Available at: https://www.medrxiv.org/content/10.1101/2024.01.09.24301030v2 [Accessed April 8, 2024].

Dutcher EG, Lopez-Cruz L, Pama EAC, Lynall M-E, Bevers ICR, Jones JA, Khan S, Sawiak SJ, Milton AL, Clatworthy MR, Robbins TW, Bullmore ET, Dalley JW (2023) Early-life stress biases responding to negative feedback and increases amygdala volume and vulnerability to later-life stress. Transl Psychiatry 13:1–11.

Ellenbroek B, Youn J (2016) Rodent models in neuroscience research: is it a rat race? Dis Model Mech 9:1079–1087.

Fenchel D et al. (2020) Development of microstructural and morphological cortical profiles in the neonatal brain. Cereb Cortex 30:5767–5779.

Fjell AM, McEvoy L, Holland D, Dale AM, Walhovd KB (2014) What is normal in normal aging? Effects of aging, amyloid and Alzheimer’s disease on the cerebral cortex and the hippocampus. Prog Neurobiol 117:20–40.

Fornito A, Zalesky A, Breakspear M (2015) The connectomics of brain disorders. Nat Rev Neurosci 16:159–172.

Fornito A, Zalesky A, Bullmore ET eds. (2016a) Chapter 6 - Components, Cores, and Clubs. In: Fundamentals of Brain Network Analysis, pp 163–206. San Diego: Academic Press.

Fornito A, Zalesky A, Bullmore ET eds. (2016b) Chapter 5 - Centrality and Hubs. In: Fundamentals of Brain Network Analysis, pp 137–161. San Diego: Academic Press.

García-Cabezas MÁ, Hacker JL, Zikopoulos B (2023) Homology of neocortical areas in rats and primates based on cortical type analysis: an update of the Hypothesis on the Dual Origin of the Neocortex. Brain Struct Funct 228:1069–1093.

González-García N, Buimer EEL, Moreno-López L, Sallie SN, Váša F, Lim S, Romero-Garcia R, Scheuplein M, Whitaker KJ, Jones PB, Dolan RJ, NSPN consortium, Fonagy P, Goodyer I, Bullmore ET, van Harmelen A-L (2023) Resilient functioning is associated with altered structural brain network topology in adolescents exposed to childhood adversity. Dev Psychopathol 35:2253–2263.

Grossman RI, Gomori JM, Ramer KN, Lexa FJ, Schnall MD (1994) Magnetization transfer: theory and clinical applications in neuroradiology. Radiographics 14:279–290.

Hahn JD, Duckworth C (2023) A brain flatmap data visualization tool for mouse, rat, and human. J Comp Neurol 531:1008–1016.

Hahn JD, Swanson LW, Bowman I, Foster NN, Zingg B, Bienkowski MS, Hintiryan H, Dong H-W (2021) An open access mouse brain flatmap and upgraded rat and human brain flatmaps based on current reference atlases. J Comp Neurol 529:576–594.

Hamano K, Takeya T, Iwasaki N, Nakayama J, Ohto T, Okada Y (1998) A quantitative study of the progress of myelination in the rat central nervous system, using the immunohistochemical method for proteolipid protein. Brain Res Dev Brain Res 108:287– 293.

Han W, Pan Y, Han Z, Cheng L, Jiang L (2022) Advanced maternal age impairs myelination in offspring rats. Front Pediatr 10:850213.

Herrick CJ (1948) The brain of the tiger salamander, Ambystoma tigrinum. Chicago: Univ. of Chicago Press.

Hettwer MD, Dorfschmidt L, Puhlmann LMC, Jacob LM, Paquola C, Bethlehem RAI, NSPN Consortium, Bullmore ET, Eickhoff SB, Valk SL (2024) Longitudinal variation in resilient psychosocial functioning is associated with ongoing cortical myelination and functional reorganization during adolescence. Nat Commun 15:6283.

Homan P, Argyelan M, DeRosse P, Szeszko PR, Gallego JA, Hanna L, Robinson DG, Kane JM, Lencz T, Malhotra AK (2019) Structural similarity networks predict clinical outcome in early-phase psychosis. Neuropsychopharmacology 44:915–922.

Jones JA, Belin-Rauscent A, Jupp B, Fouyssac M, Sawiak SJ, Zuhlsdorff K, Zhukovsky P, Hebdon L, Velazquez Sanchez C, Robbins TW, Everitt BJ, Belin D, Dalley JW (2024) Neurobehavioral Precursors of Compulsive Cocaine Seeking in Dual Frontostriatal Circuits. Biological Psychiatry Global Open Science 4:194–202.

Jung B, Taylor PA, Seidlitz J, Sponheim C, Perkins P, Ungerleider LG, Glen D, Messinger A (2021) A comprehensive macaque fMRI pipeline and hierarchical atlas. Neuroimage 235:117997.

Kleven H, Bjerke IE, Clascá F, Groenewegen HJ, Bjaalie JG, Leergaard TB (2023) Waxholm Space atlas of the rat brain: a 3D atlas supporting data analysis and integration. Nat Methods 20:1822–1829.

Kolk SM, Rakic P (2022) Development of prefrontal cortex. Neuropsychopharmacology 47:41– 57.

Krigman MR, Hogan EL (1976) Undernutrition in the developing rat: effect upon myelination. Brain Res 107:239–255.

Lein ES et al. (2007) Genome-wide atlas of gene expression in the adult mouse brain. Nature 445:168–176.

Lerch JP, Yiu AP, Martinez-Canabal A, Pekar T, Bohbot VD, Frankland PW, Henkelman RM, Josselyn SA, Sled JG (2011) Maze training in mice induces MRI-detectable brain shape changes specific to the type of learning. Neuroimage 54:2086–2095.

Li W, Yang C, Shi F, Wu S, Wang Q, Nie Y, Zhang X (2017) Construction of individual morphological brain networks with multiple morphometric features. Front Neuroanat 11:34.

Li X, Wu K, Zhang Y, Kong L, Bertisch H, DeLisi LE (2019) Altered topological characteristics of morphological brain network relate to language impairment in high genetic risk subjects and schizophrenia patients. Schizophr Res 208:338–343.

Lisowska A, Rekik I (2019) Joint pairing and structured mapping of convolutional brain morphological multiplexes for early dementia diagnosis. Brain Connect 9:22–36.

Long KLP, Chao LL, Kazama Y, An A, Hu KY, Peretz L, Muller DCY, Roan VD, Misra R, Toth CE, Breton JM, Casazza W, Mostafavi S, Huber BR, Woodward SH, Neylan TC, Kaufer D (2021) Regional gray matter oligodendrocyte- and myelin-related measures are associated with differential susceptibility to stress-induced behavior in rats and humans. Transl Psychiatry 11:1–15.

Lupien SJ, McEwen BS, Gunnar MR, Heim C (2009) Effects of stress throughout the lifespan on the brain, behaviour and cognition. Nat Rev Neurosci 10:434–445.

Mahjoub I, Mahjoub MA, Rekik I, Alzheimer’s Disease Neuroimaging Initiative (2018) Brain multiplexes reveal morphological connectional biomarkers fingerprinting late brain dementia states. Sci Rep 8:4103.

Mancini M, Karakuzu A, Cohen-Adad J, Cercignani M, Nichols TE, Stikov N (2020) An interactive meta-analysis of MRI biomarkers of myelin. Elife 9 Available at: https://elifesciences.org/articles/61523 [Accessed September 26, 2024].

Meier S, Bräuer AU, Heimrich B, Nitsch R, Savaskan NE (2004) Myelination in the hippocampus during development and following lesion. Cell Mol Life Sci 61:1082–1094.

Mengler L, Khmelinskii A, Diedenhofen M, Po C, Staring M, Lelieveldt BPF, Hoehn M (2014) Brain maturation of the adolescent rat cortex and striatum: changes in volume and myelination. Neuroimage 84:35–44.

Morgan SE, Seidlitz J, Whitaker KJ, Romero-Garcia R, Clifton NE, Scarpazza C, van Amelsvoort T, Marcelis M, van Os J, Donohoe G, Mothersill D, Corvin A, Pocklington A, Raznahan A, McGuire P, Vértes PE, Bullmore ET (2019) Cortical patterning of abnormal morphometric similarity in psychosis is associated with brain expression of schizophrenia-related genes. Proc Natl Acad Sci U S A 116:9604–9609.

Nauta WJH, Karten HJ (1970) A General Profile of the Vertebrate Brain, with Sidelights on the Ancestry of Cerebral Cortex. In: Neuroanatomy (Nauta WJH, ed), pp 520–539. Boston, MA: Birkhäuser.

Ohline SM, Abraham WC (2019) Environmental enrichment effects on synaptic and cellular physiology of hippocampal neurons. Neuropharmacology 145:3–12.

Paus T, Keshavan M, Giedd JN (2008) Why do many psychiatric disorders emerge during adolescence? Nat Rev Neurosci 9:947–957.

Paxinos G, Watson C (2006) The Rat Brain in Stereotaxic Coordinates: Hard Cover Edition. Elsevier.

Pebesma E (2018) Simple features for R: Standardized support for spatial vector data. R J 10:439.

Roussel J-R, Auty D, Coops NC, Tompalski P, Goodbody TRH, Meador AS, Bourdon J-F, de Boissieu F, Achim A (2020) lidR: An R package for analysis of Airborne Laser Scanning (ALS) data. Remote Sens Environ 251:112061.

Ruan J, Wang N, Li J, Wang J, Zou Q, Lv Y, Zhang H, Wang J (2023) Single-subject cortical morphological brain networks across the adult lifespan. Hum Brain Mapp 44:5429–5449.

Rubinov M, Ypma RJ, Watson C, Bullmore ET (2015) Wiring cost and topological participation of the mouse brain connectome. Proceedings of the National Academy of Sciences 112:10032–10037.

Sánchez MM, Ladd CO, Plotsky PM (2001) Early adverse experience as a developmental risk factor for later psychopathology: evidence from rodent and primate models. Dev Psychopathol 13:419–449.

Sebenius I, Dorfschmidt L, Seidlitz J, Alexander-Bloch A, Morgan SE, Bullmore E (2024) Structural MRI of brain similarity networks. Nat Rev Neurosci:1–18.

Sebenius I, Seidlitz J, Warrier V, Bethlehem RAI, Alexander-Bloch A, Mallard TT, Garcia RR, Bullmore ET, Morgan SE (2023) Robust estimation of cortical similarity networks from brain MRI. Nat Neurosci 26:1461–1471.

Seidlitz J et al. (2020) Transcriptomic and cellular decoding of regional brain vulnerability to neurogenetic disorders. Nat Commun 11:3358.

Seidlitz J, Váša F, Shinn M, Romero-Garcia R, Whitaker KJ, Vértes PE, Wagstyl K, Kirkpatrick Reardon P, Clasen L, Liu S, Messinger A, Leopold DA, Fonagy P, Dolan RJ, Jones PB, Goodyer IM, Raznahan A, Bullmore ET (2018) Morphometric Similarity Networks Detect Microscale Cortical Organization and Predict Inter-Individual Cognitive Variation. Neuron 97:231–247.e7.

Stein LR, O’Dell KA, Funatsu M, Zorumski CF, Izumi Y (2016) Short-term environmental enrichment enhances synaptic plasticity in hippocampal slices from aged rats. Neuroscience 329:294–305.

Swanson LW (1992) Brain maps: structure of the rat brain, 1st ed.

Swanson LW (2000) 3 - A History of Neuroanatomical Mapping. In: Brain Mapping: The Systems (Toga AW, Mazziotta JC, eds), pp 77–109. San Diego: Academic Press.

Swanson LW (2018) Brain maps 4.0—Structure of the rat brain: An open access atlas with global nervous system nomenclature ontology and flatmaps. J Comp Neurol 526:935– 943.

Swanson LW, Hahn JD, Sporns O (2017) Organizing principles for the cerebral cortex network of commissural and association connections. Proceedings of the National Academy of Sciences 114:E9692–E9701.

Swanson LW, Hahn JD, Sporns O (2020) Structure–function subsystem models of female and male forebrain networks integrating cognition, affect, behavior, and bodily functions. Proceedings of the National Academy of Sciences 117:31470–31481.

Swanson LW, Hahn JD, Sporns O (2022) Structure–function subsystem model and computational lesions of the central nervous system’s rostral sector (forebrain and midbrain). Proceedings of the National Academy of Sciences 119:e2210931119.

Swanson LW, Hahn JD, Sporns O (2024a) Network architecture of intrinsic connectivity in a mammalian spinal cord (the central nervous system’s caudal sector). Proc Natl Acad Sci U S A 121:e2320953121.

Swanson LW, Hahn JD, Sporns O (2024b) Neural network architecture of a mammalian brain. Proc Natl Acad Sci U S A 121:e2413422121.

Swanson LW, Sporns O, Hahn JD (2019) The network architecture of rat intrinsic interbrain (diencephalic) macroconnections. Proc Natl Acad Sci U S A 116:26991–27000.

Tian T, Li J, Zhang G, Wang J, Liu D, Wan C, Fang J, Wu D, Zhou Y, Zhu W (2021) Effects of childhood trauma experience and BDNF Val66Met polymorphism on brain plasticity relate to emotion regulation. Behav Brain Res 398:112949.

Towlson EK, Vértes PE, Ahnert SE, Schafer WR, Bullmore ET (2013) The rich club of the C. elegans neuronal connectome. J Neurosci 33:6380–6387.

van den Heuvel MP, Sporns O (2011) Rich-club organization of the human connectome. J Neurosci 31:15775–15786.

van den Heuvel MP, Sporns O (2013) Network hubs in the human brain. Trends Cogn Sci 17:683–696.

Wickham H (2016) Ggplot2: Elegant graphics for data analysis, 2nd ed. Cham, Switzerland: Springer International Publishing.

Wu X, Palaniyappan L, Yu G, Zhang K, Seidlitz J, Liu Z, Kong X, Schumann G, Feng J, Sahakian BJ, Robbins TW, Bullmore E, Zhang J (2023) Morphometric dis-similarity between cortical and subcortical areas underlies cognitive function and psychiatric symptomatology: a preadolescence study from ABCD. Mol Psychiatry 28:1146–1158.

Xiao Y, Chen F, Lei W, Ke J, Dai Y, Qi R, Lu G, Zhong Y (2023) Transcriptional signal and cell specificity of genes related to cortical structural differences of post-traumatic stress disorder. J Psychiatr Res 160:28–37.

Yao Z et al. (2023) A high-resolution transcriptomic and spatial atlas of cell types in the whole mouse brain. Nature 624:317–332.

Zeiss CJ (2021) Comparative milestones in rodent and human postnatal central nervous system development. Toxicol Pathol 49:1368–1373.

Zhang W, Lei D, Keedy SK, Ivleva EI, Eum S, Yao L, Tamminga CA, Clementz BA, Keshavan MS, Pearlson GD, Gershon ES, Bishop JR, Gong Q, Lui S, Sweeney JA (2020) Brain gray matter network organization in psychotic disorders. Neuropsychopharmacology 45:666–674.

Zilles K (2012) The Cortex of the Rat: A Stereotaxic Atlas. Springer Science & Business Media.

