## Supplemental material for "The brain cortical similarity network: Development and sensitivity to early life stress in a rat model"

### This PDF file includes

Figures S1 to S5
Legends for Tables S1 and S2
SI References

### Other supporting materials for this manuscript include the following

Tables S1 and S2

**Figure S1.**

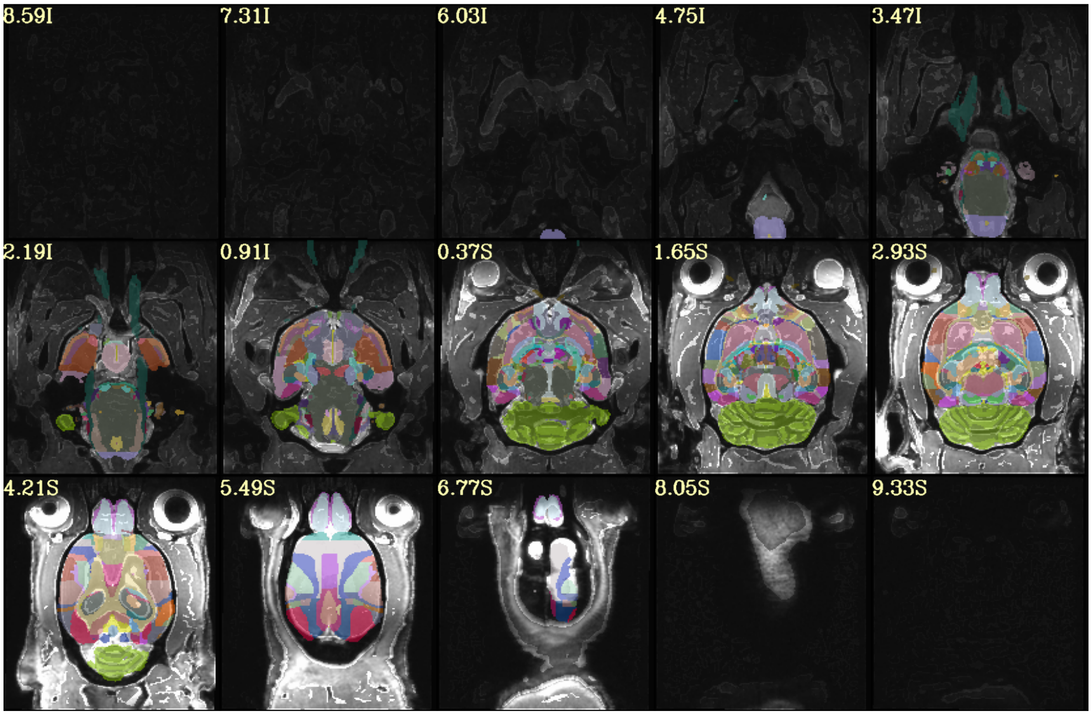


**Example quality control images in the axial plane;** direct output from the AFNI @animal_warper function. The pipeline also provides quality control (QC) images in the coronal and sagittal slices. The example subject is a post-natal day (PND) 63 rat from the normative developmental cohort.

**Figure S2.**

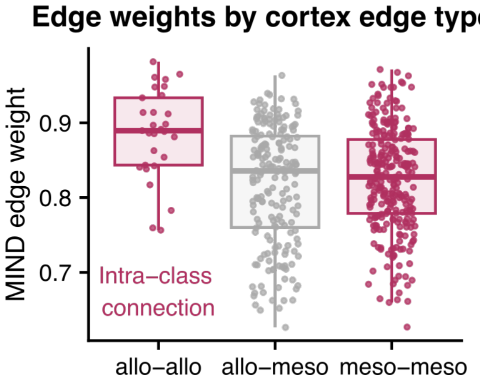


**Distributions of intra-cortex class and inter-cortex class morphometric inverse divergence (MIND) edge weights**. Maroon indicates a within cortex type connection, while gray indicates a between cortex type connection (allo = allocortex, meso = mesocortex). Eulaminate (isocortical) areas were not considered in this analysis due to too few representative regions.

**Figure S3.**

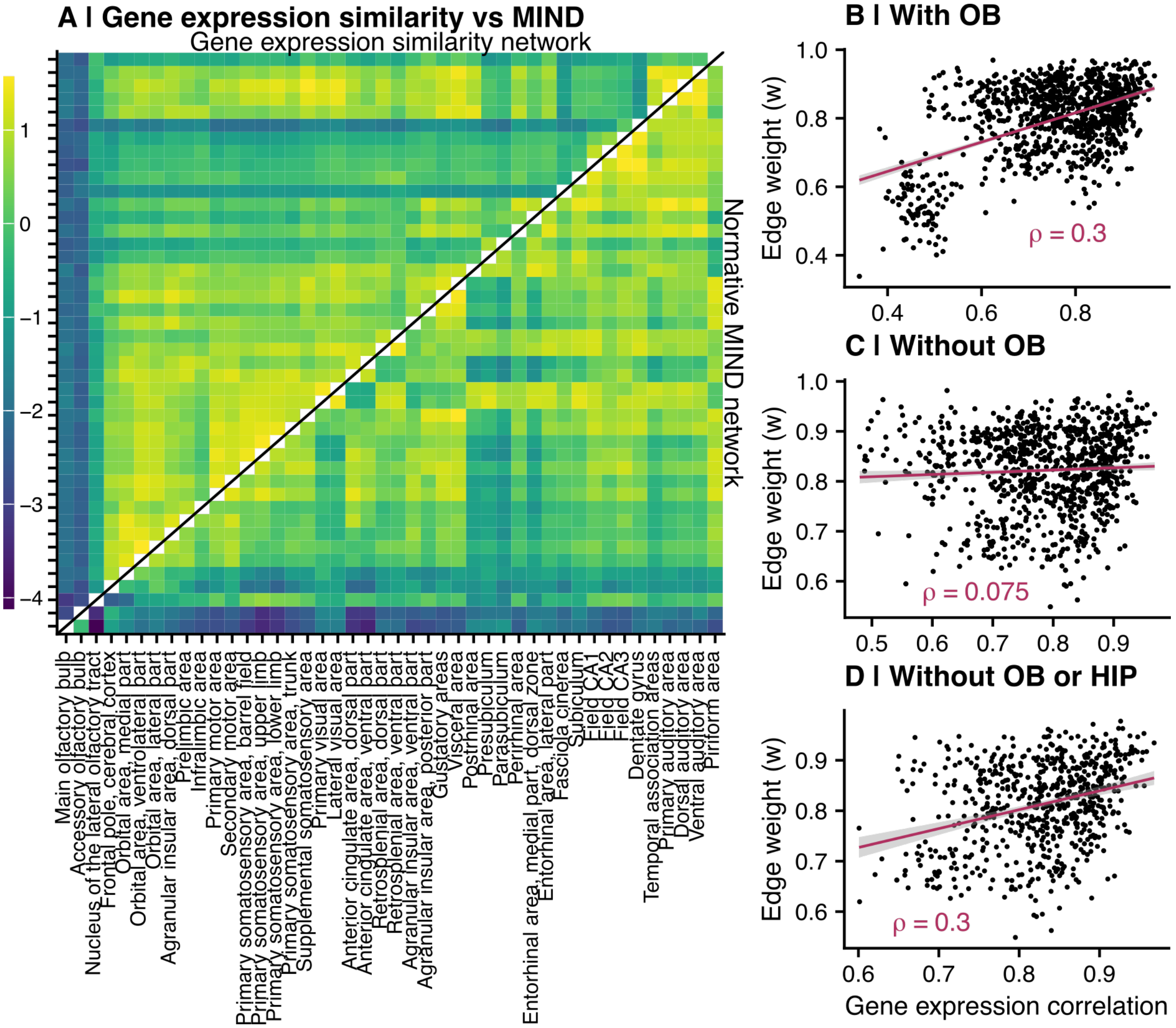


**The normative MIND network aligns with spatial gene expression similarity defined in the mouse brain. A)** Heatmap representations of the mouse gene expression similarity network (top left of the diagonal) and weighted MIND similarity network (bottom right of the diagonal). Both networks include olfactory bulb (OB) regions in this visualization. Each row and column represent a region of interest, defined by the AMBA (Lein et al., 2007). To increase comparability between the networks, weights were *Z*-scored. **B)** Correlation between mouse gene expression similarity (x-axis) and normative MIND edge weight (y-axis) when olfactory bulb (OB) regions are included. Each point represents an edge; the line of best fit and Spearman correlation are shown in maroon. **C)** Same as panel B but without any OB regions. **D)** Edges with low similarity in mouse gene expression profiles, but high MIND similarity, principally comprising hippocampal regions, with removal of these edges improving the correlation between MIND and gene expression similarity (ρ=0.32).

**Figure S4.**

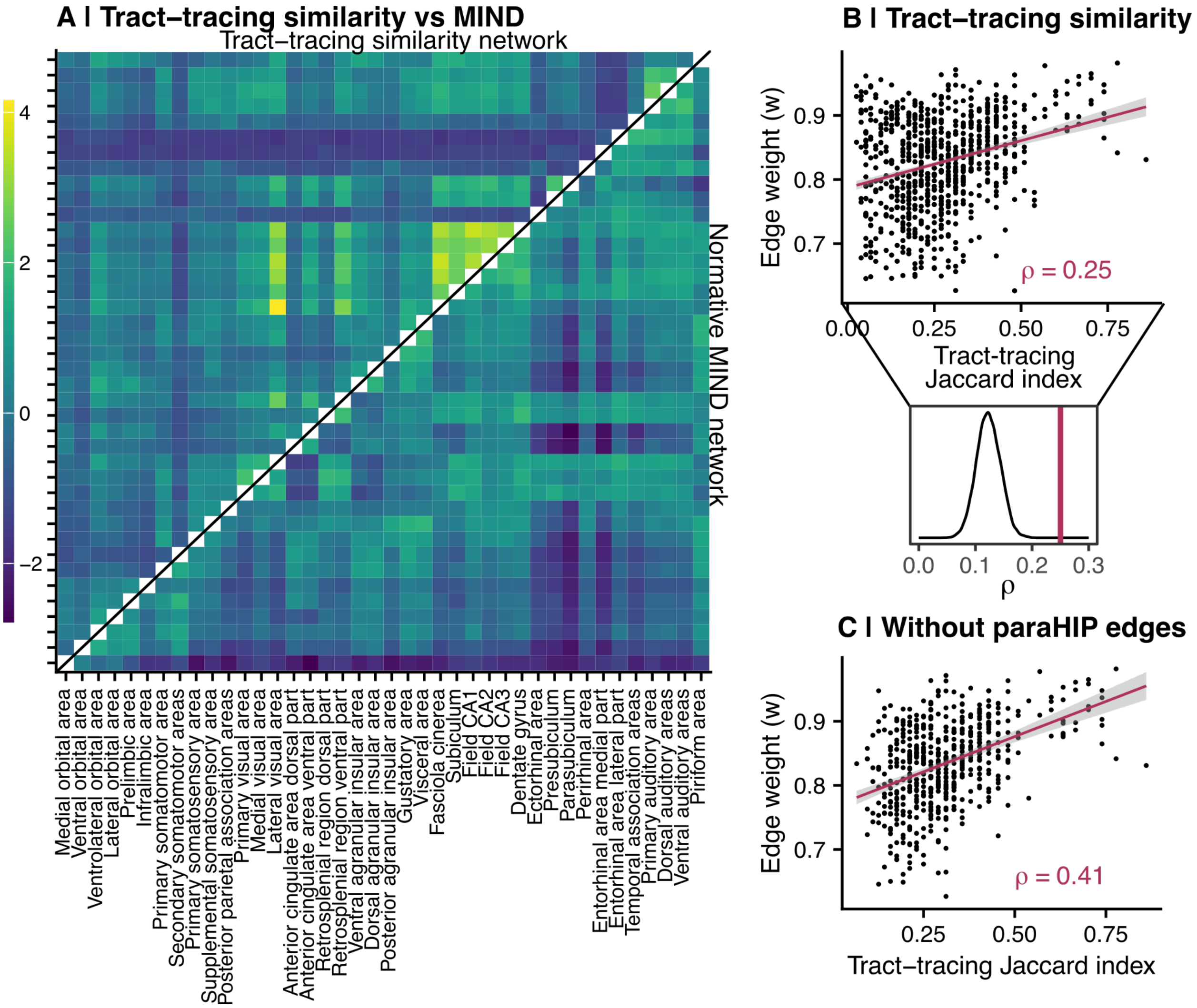


**The normative MIND network reflects similarity of axonal connectivity. A)** Heatmap representations of the tract-tracing similarity network (top left of the diagonal) and weighted MIND similarity network (bottom right of the diagonal). Each row and column represent a region of interest, defined by the Brain Maps 4.0 atlas (Swanson, 2018). To increase comparability between the networks, weights were *Z*-scored. **B)** Top: Correlation between similarity of tract-tracing connection profiles between pairwise combinations of regions and strength of MIND similarity (same as **Figure 4D**). Bottom: This relationship was significant compared to a null distribution of 10000 distance-corrected networks (normative network Spearman correlation $Z$=6; $P$<0.001). **C)** Edges with low similarity in tract-tracing profiles, but high MIND similarity, principally comprised parahippocampal regions, with removal of these edges improving the correlation between MIND and tract-tracing similarity (ρ=0.41).

**Figure S5.**

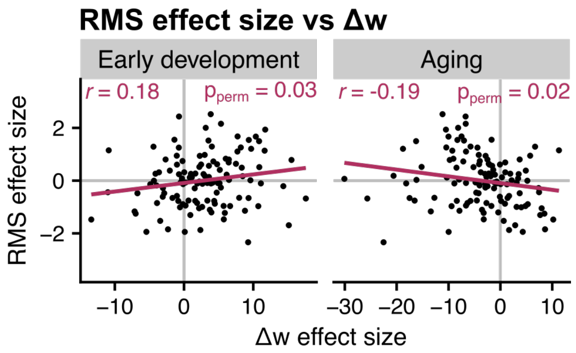


**The relationship between normative developmental change and repeated maternal separation (RMS)-induced network perturbations in early adulthood.** The x-axis represents Δw from the normative developmental cohort (left = early development; right = aging), while the y-axis shows the PND 63 RMS-control effect size from the experimental cohort. Each point represents a system-level edge, while the line of best fit is shown in maroon. RMS edge assignments were permuted 10000 times and the actual Pearson correlation was scaled in relation to the null distribution. RMS effect size was significantly associated with both $\Delta w_{dev}$ ($r$=0.18, $P_{perm}$=0.03, $Z_{perm}$=1.90) and $\Delta w_{age}$ ($r$=-0.19, $P_{perm}$=0.02, $Z_{perm}$=-2.03).

### Legends for Supplemental Tables S1 and S2

**Table S1 (separate file).** The {53×53} matrix representation of the normative cortical MIND network, defined as the median edge weight across N=41 individuals at the PND 63 timepoint in the normative developmental cohort.

**Table S2 (separate file).** Anatomical correspondence of the Waxholm Space Atlas to other rat brain atlases, including:

A. Brain Maps 4 (Swanson, 2018)
B. Zilles atlas (Zilles, 2012)
C. Allen Mouse Brain Atlas (Lein et al., 2007)

Atlas mappings were generated using the methodology described in the **Methods** section.

**SI References**

Lein ES et al. (2007) Genome-wide atlas of gene expression in the adult mouse brain. Nature 445:168–176.

Swanson LW (2018) Brain maps 4.0—Structure of the rat brain: An open access atlas with global nervous system nomenclature ontology and flatmaps. J Comp Neurol 526:935–943.

Zilles K (2012) The Cortex of the Rat: A Stereotaxic Atlas. Springer Science & Business Media.
